# Production of moth sex pheromones for pest control by yeast fermentation

**DOI:** 10.1101/2020.07.15.205047

**Authors:** Carina Holkenbrink, Bao-Jian Ding, Hong-Lei Wang, Marie Inger Dam, Karolis Petkevicius, Kanchana Rueksomtawin Kildegaard, Leonie Wenning, Christina Sinkwitz, Bettina Lorántfy, Eleni Koutsoumpeli, Lucas França, Marina Pires, Carmem Bernardi, William Urrutia, Agenor Mafra-Neto, Bruno Sommer Ferreira, Dimitris Raptopoulos, Maria Konstantopoulou, Christer Löfstedt, Irina Borodina

**Affiliations:** The Novo Nordisk Foundation Center for Biosustainability, Technical University of Denmark, Kemitorvet 220, 2800 Kgs. Lyngby, Denmark; BioPhero ApS, Kemitorvet 378, 2800 Kgs. Lyngby, Denmark; Lund University, Department of Biology, Sölvegatan 37, SE-224 60 Lund, Sweden; Chemical Ecology and Natural Products Laboratory, Institute of Biosciences and Applications, National Centre of Scientific Research Demokritos, Attikis, Greece; Biotrend S.A., Biocant Park, Núcleo 04 Lote 2, 3060-197 Cantanhede, Portugal; ISCA Technologies, 1230 W. Spring St. Riverside, California 92507, USA; Novagrica Hellas S.A., TESPA “Lefkippos”, 15341, Athens, Greece

**Author notes:** Correspondence and requests for materials should be addressed to Irina Borodina or Christer Löfstedt.

## Abstract

The use of insect sex pheromones is an alternative technology for pest control in agriculture and forestry, which, in contrast to insecticides, does not have adverse effects on human health or environment and is efficient also against insecticide-resistant insect populations.^1,2^ Due to the high cost of chemically synthesized pheromones, mating disruption applications are currently primarily targeting higher value crops, such as fruits.^3^ Here we demonstrate a biotechnological method for the production of pheromones of economically important moth pests using engineered yeast cell factories. Biosynthetic pathways towards several pheromones or their precursors were reconstructed in the oleaginous yeast *Yarrowia lipolytica*, which was further metabolically engineered for improved pheromone biosynthesis by decreasing fatty alcohol degradation and downregulating storage lipid accumulation. The sex pheromone of the cotton bollworm *Helicoverpa armigera* was produced by oxidation of fermented fatty alcohols into corresponding aldehydes. The resulting pheromone was just as efficient and specific for trapping of *H. armigera* male moths in cotton fields in Greece as a synthetic pheromone mixture. We further demonstrated the production of the main pheromone component of the fall armyworm *Spodoptera frugiperda*. Our work describes a biotech platform for the production of commercially relevant titres of moth pheromones for pest control by yeast fermentation.

**Significance statement:** Agriculture largely relies on insecticides and genetically modified crops for pest control, however alternative solutions are required due to emerging resistance, toxicity and regulatory issues, and consumer preferences. Mating disruption with sex pheromones that act by preventing insect reproduction is considered the most promising and scalable alternative to insecticides. This method is highly efficient and safe for human health and environment. The likelihood of insect resistance development is very low and can be handled by adjusting the pheromone composition. The high cost of chemically synthesized pheromones is the major barrier for the wider adoption of pheromones. A novel method based on yeast fermentation enables the production of insect sex pheromones as a lower cost from renewable feedstocks.

## MAIN TEXT

### Establishing pathways towards moth pheromones in yeast

The majority of identified moth sex pheromone components are unsaturated acetates, alcohols or aldehydes. These compounds are derived from the fatty acid metabolism and possess 10-18 carbon atoms in the carbon skeleton. They are called Type I moth pheromone components and constitute approximately 75% of all known moth sex pheromone components.^4,5^ The chemical diversity is to a large extent produced by the combined action of specific desaturases and limited chain-shortening^6^. To establish pathways towards moth pheromone compounds in yeast, we first investigated a range of fatty acyl-CoA desaturases and reductases for the production of (*Z*)-hexadec-11-en-1-ol (*Z*11-16OH) (Fig. 1A) by expressing the enzymes in combinations in baker’s yeast *Saccharomyces cerevisiae*. The fermented *Z*11-16OH can be chemically oxidized into (*Z*)-hexadec-1 1 -enal (*Z*11-16Ald), which is the main sex pheromone component of several row crop pests, such as the cotton bollworm *Helicoverpa armigera*, the striped rice stemborer *Chilo suppressalis*, and the yellow rice stemborer *Scirpophaga incertulas*^7^. The combination of a desaturase from *Amyelois transitella* and a reductase from *H*. *armigera* resulted in the highest titre of 1.7±0.4 mg/L *Z*11-16OH (Fig. 1B), which was an order of magnitude enhancement in comparison to the previous study.^8^ The improvement was likely due to the utilization of a desaturase variant with a higher activity in yeast and due to expression of the genes from constitutive promoters using constructs stably integrated into the yeast genome.^9^ Next, we wanted to achieve the biosynthesis of (*Z*)-tetradec-9-en-1-yl acetate (*Z*9-14Ac), which is the main sex pheromone component of the fall armyworm *Spodoptera frugiperda*, a rising pest with a high occurrence of insecticide resistance.^10,11^ For this, we searched for a Δ9-desaturase with a higher activity and specificity towards tetradecanoyl-CoA than to hexadecanoyl-CoA (Fig. 1C). The activities of six heterologous desaturase candidates were tested in a *S. cerevisiae ole1Δelo1Δ* strain devoid of the native desaturation and elongation activities. The cells were cultivated with supplementation of methyl tetradecanoate (14Me) and the total lipids were analysed to determine the desaturated fatty acids (Fig. 1D). The strain expressing the desaturase from *Drosophila melanogaster* resulted in the highest concentration of 3.67±0.99 mg/L methyl (*Z*)-tetradec-9-enoate (*Z*9-14Me) and in the highest *Z*9-14Me to methyl (*Z*)-hexadec-9-enoate (*Z*9-16Me) ratio, indicating a higher specificity towards the tetradecanoyl-CoA substrate. To establish the complete pathway towards *Z*9-14Ac in the yeast *S. cerevisiae*, we expressed the *D. melanogaster* Δ9 desaturase together with the *H. armigera* reductase and *S. cerevisiae ATF1* known to catalyse acetylation of fatty alcohols^12^. The resulting strain produced 7.3±0.2 mg/L of *Z*9-14Ac in comparison to 1.4±0.4 mg/L in an analogous strain lacking the heterologous Δ9 desaturase (Fig. 1E).

**Fig. 1.**
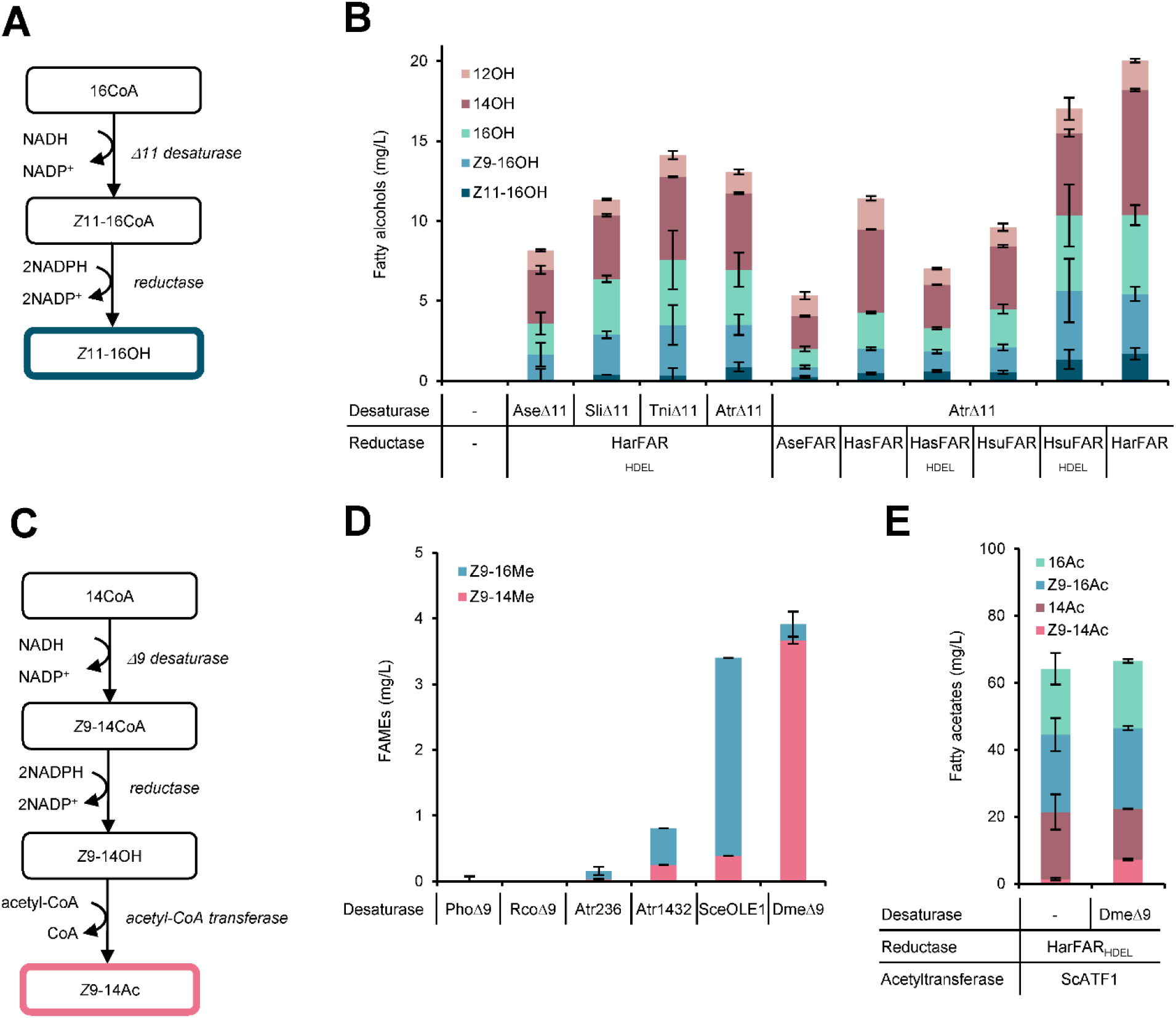
Biosynthesis of moth sex pheromone compounds in yeast. **A.** Biosynthetic pathway towards (*Z*)-hexadec-11-en-1-ol (*Z*11-16OH). **B.** Concentrations of fatty alcohols in the cultures of *S. cerevisiae* expressing combinations of desaturases and reductases from different lepidopteran species. **C**. Biosynthetic pathway towards (*Z*)-tetradec-9-en-1-yl acetate (*Z*9-14Ac). **D.** Concentrations of fatty acids (analysed as methyl esters) in *S. cerevisiae* cells that express plant and insect desaturases. **E.** Concentrations of fatty alcohol acetates in cultures of *S. cerevisiae* expressing combinations of desaturase, reductase and acetyl-CoA transferase genes. The cultivations were performed at small-scale in tubes or deep-well plates in biological triplicates. The average values and standard deviations are shown. Abbreviations: AseΔ11 - Δ11 desaturase from *Agrotis segetum*, SliΔ11 desaturase - Δ11 desaturase from *Spodoptera littoralis*, TniΔ11 - Δ11 desaturase from *Trichoplusia ni*, AtrΔ11 - Δ11 desaturase from *Amyelois transitella*, HarFAR – fatty acyl-CoA reductase from *Helicoverpa armigera*, AseFAR – fatty acyl-CoA reductase from *A. segetum*, HasFAR – fatty acyl-CoA reductase from *Helicoverpa assulta*, HsuFAR – fatty acyl-CoA reductase from *Heliothis subflexa*, PhoΔ9 - Δ9 desaturase from *Pelargonium x hortorum*, RcoΔ9 - Δ9 desaturase from *Ricinus communis*, Atr236 – desaturase 236 from *A. transitella*, Atr1432 – desaturase 1432 from *A. transitella*, SceOLE1 - Δ9 desaturase from *S. cerevisiae*, DmeΔ9 - Δ9 desaturase from *Drosophila melanogaster*, SceATF1 - alcohol acetyltransferase from *S. cerevisiae*, FAR_HDEL_ – modified desaturase, where the C-terminal insect signal peptide was replaced with 4-amino acid signal peptide from *S. cerevisiae*.

### Optimization of the oleaginous yeast *Yarrowia lipolytica* for moth pheromone production

We rationalized that an oleaginous yeast should be a more suitable cell factory for production of fatty alcohol-based moth pheromones than the baker’s yeast that has a low content of the fatty acid precursor acetyl-CoA in the cytosol and can only accumulate small amounts of intracellular lipids. In contrast, the oleaginous yeast *Yarrowia lipolytica* has a naturally high fatty acid metabolism and has been engineered for commercial production of polyunsaturated omega-3 fatty acids^13^ and for production of lipids^14^. Robust genetic tools, including the CRISPR/Cas9 method, have recently been developed for *Y. lipolytica*, and allow the rapid engineering of this yeast species.^15,16^ The first hurdle we encountered when co-opting *Y. lipolytica* for the production of pheromones, was the prevention of endogenous degradation of the target fatty alcohols *Z*11-16OH and (*Z*)-tetradec-9-en-1-ol (*Z*9-14OH). We deleted one by one and in combination the genes encoding the enzymes potentially implicated in fatty alcohol degradation: fatty aldehyde dehydrogenases Hfd1p and Hfd4p^17^, as well as fatty alcohol oxidase Fao1p^18^ (Fig. 2A). Moreover, we deleted peroxisomal biogenesis factor Pex10p, thus interrupting the correct assembly of peroxisomes and preventing acyl-CoA degradation. Single deletions of *HFD1/HFD4/FAO1/PEX10* genes increased the titre of *Z*11-16OH two- to three-fold, while the combination of four deletions resulted in a 19-fold titre increase (Fig. 2B). The quadruple deletion strain (ST5789) produced 14.9±3.6 mg/L of *Z*11-16OH in comparison to 0.8±0.1 mg/L produced by a reference strain only expressing the biosynthetic pathway towards *Z*11-16OH (ST3844). When strains ST3844 and ST5789 were incubated with externally supplied 1 g/L *Z*11-16OH and 1 g/L *Z*9-14OH each, strain ST3844 largely degraded the supplied alcohols, with only 6.2±3.8 mg/L *Z*11-16OH left at the end of the cultivation. In contrast to that, strain ST5789 showed a remaining concentration of 630.9±137.1 mg/L *Z*9-14OH and 620.3±73.9 mg/L *Z*11-16OH. Less than 1 g/L of fatty alcohols were recovered probably due to evaporation and some losses during the recovery procedure. A control, which contained only cultivation medium and externally supplied fatty alcohols, showed a remaining concentration of 500.7±135.7 mg/L *Z*9-14OH and 536.9±166.1 mg/L *Z*11-16OH (Fig. S1). The experiment confirmed that the degradation rate of fatty alcohols was much lower in the strain with deletion of *HFD1, HFD4, PEX10*, and *FAO1* genes.

**Fig. 2.**
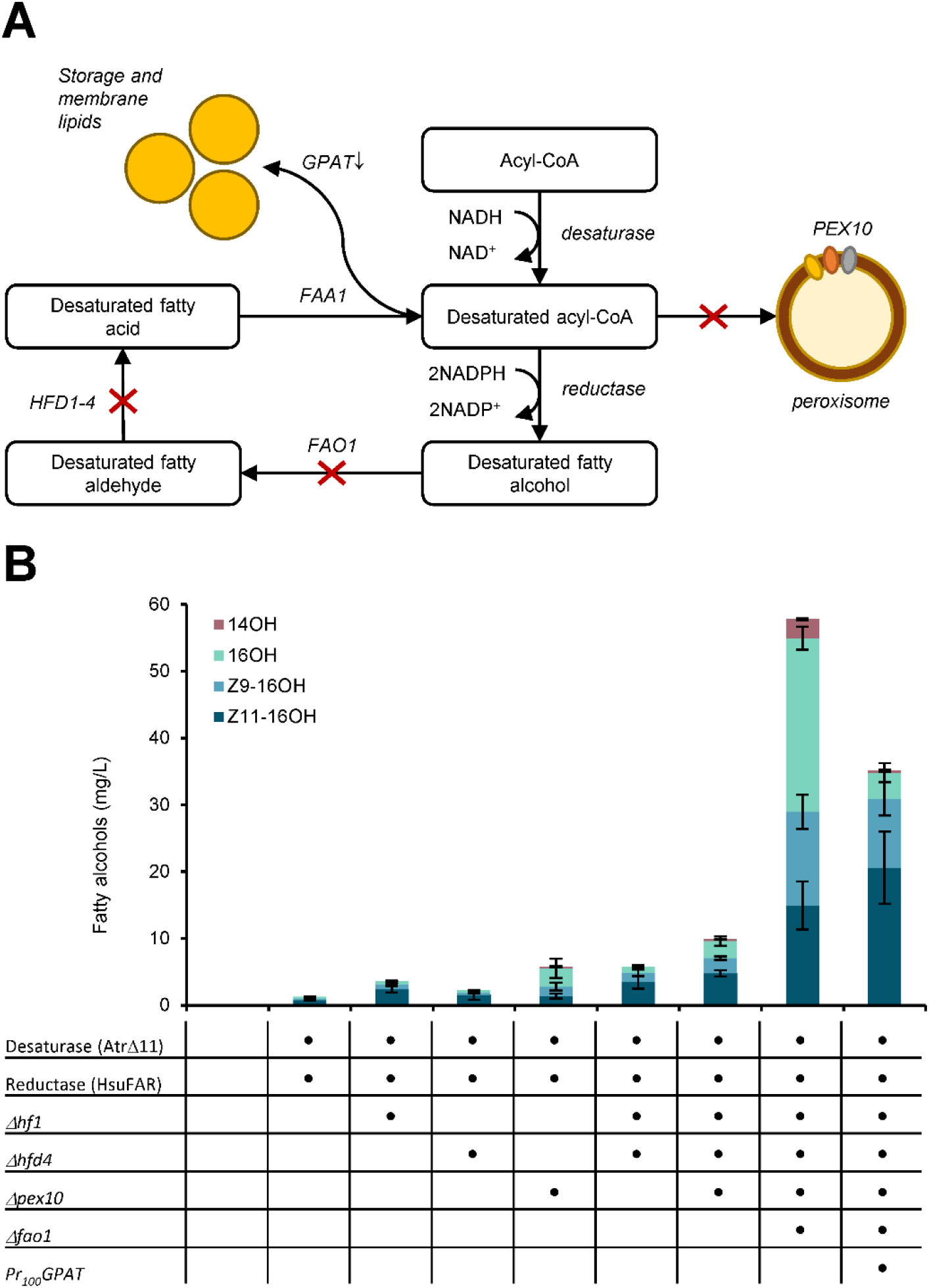
Metabolic engineering of the oleaginous yeast *Y. lipolytica* towards enhanced production of moth pheromone compounds. **A.** Overview of metabolic engineering strategies. **B.** Concentrations of fatty alcohols produced by engineered *Y. lipolytica* strains. The cultivations were performed at small-scale in tubes or deep-well plates in biological triplicates. The average values and standard deviations are shown. Abbreviations: *PEX10* – peroxisomal biogenesis factor, *FAO1* – fatty alcohol oxidase, *HFD1-4* – fatty aldehyde dehydrogenases, *FAA1* – fatty acyl-CoA synthetase, *GPAT* – glycerol-3-phosphate acyltransferase.

Another challenge with *Y. lipolytica* as a host was to reduce the channelling of fatty acyl-CoAs, the fatty alcohol precursors, into storage lipids. We hence downregulated the expression of the gene encoding glycerol-3-phosphate acyltransferase (GPAT), which catalyses the first committing step towards glycerolipid- and glycerophospholipid biosynthesis. The downregulation was achieved by truncating the *GPAT* promoter to 100 base pairs and confirmed by qRT-PCR (Fig. S2.A). The downregulation of *GPAT* improved the titre of *Z*11-16OH from 14.9±3.6 mg/L to 20.6±5.4 mg/L (Fig. 2B). At the same time, the total fatty acid content of the cells was reduced by 53% (Fig. S2.B, C). The combination of *Y. lipolytica* genome edits that reduce the fatty alcohol degradation and lipid accumulation thus resulted in a basic platform chassis, where various moth pheromone pathways can be inserted.

The strain, however, predominantly produced fatty alcohols of 16-carbon chain length. In order to enable the production of 14-carbon pheromones, we introduced a mutation into fatty acid synthase subunit Fas2p^I1220F^, which was previously reported to benefit the biosynthesis of tetradecanoyl-CoA.^19^ We expressed the pathway towards *Z*9-14OH in the engineered *Y. lipolytica* strains and obtained 4.9±1.4 mg/L titre in the basic platform chassis and 73.6±16 mg/L *Z*9-14OH in the platform chassis with additional Fas2p mutation (Fig. 3B). The mutation thus resulted in a 15-fold improvement of a 14-carbon product and should be beneficial for producing also other pheromones derived from tetradecanoyl-CoA.

**Fig. 3.**
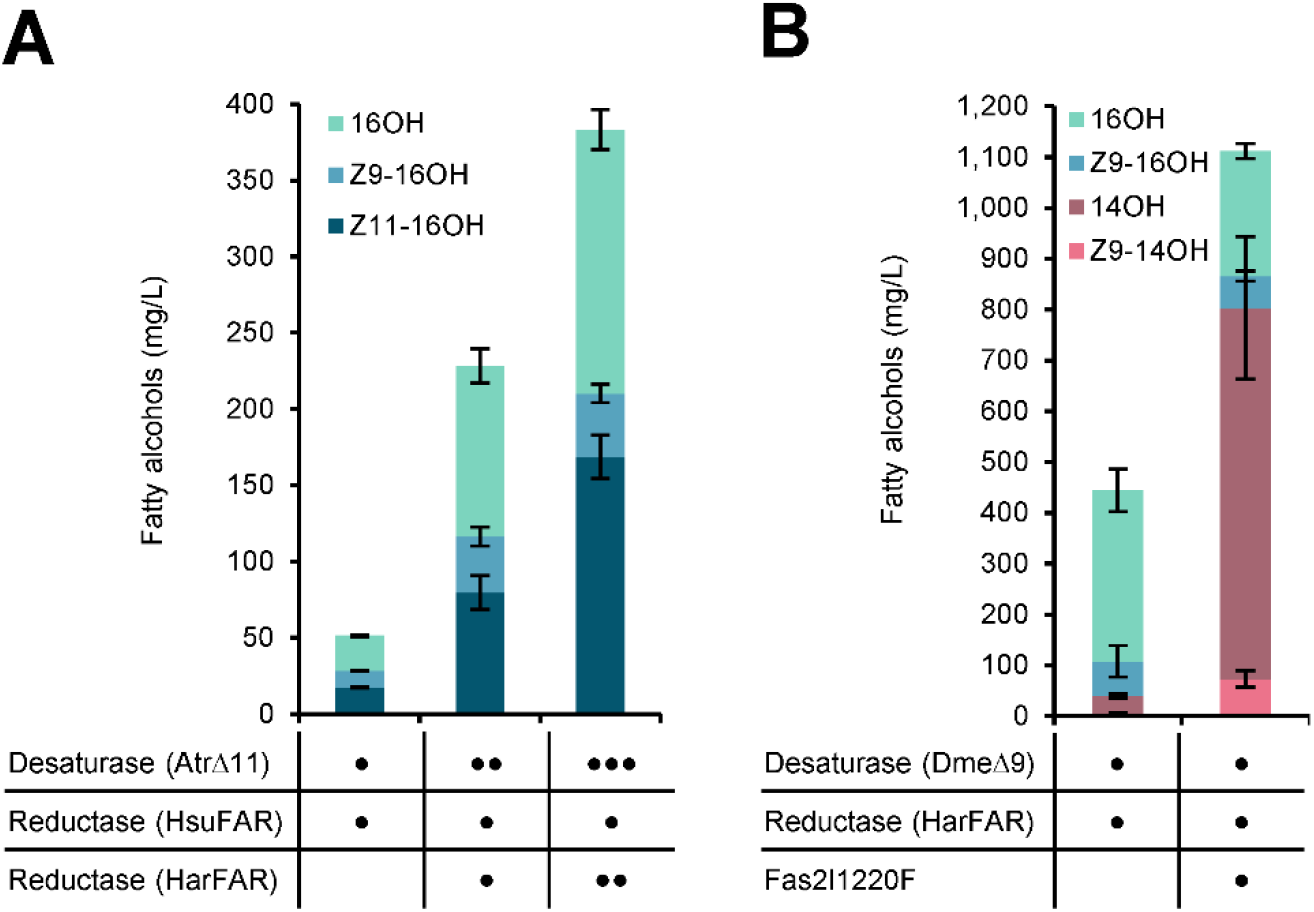
Production of two lepidopteran pheromone compounds in engineered *Y. lipolytica* strains. **A.** Production of (*Z*)-hexadec-11-en-1-ol (*Z*11-16OH) in the engineered *Y. lipolytica* host strain (Δ*hfd1Δhfd4Δpex10Δfao1*P_GPAT_100) harbouring different copy numbers of the pathway. The dots indicate the number of the integrated gene copies. **B.** Production of (*Z*)-tetradec-9-en-1-ol (*Z*9-14OH) in the *Y. lipolytica* host strain additionally engineered by mutating the fatty acid synthase subunit *FAS2*. The cultivations were performed at small-scale in tubes or deep-well plates in biological triplicates. The average values and standard deviations are shown. The gene name abbreviations are as in Fig. 1.

To further improve the production of *Z*11-16OH, we integrated additional copies of desaturase and reductase genes to pull the flux towards pheromone biosynthesis. Integration of the second copy of the pathway increased the titre 4.6-fold. The strain with three copies of the pathway produced 169±14 mg/L *Z*11-16OH, a 9.7-fold increase in comparison to the single copy strain (Fig 3A). When the optimized yeast strain was fermented in a 10L-bioreactor, 2.57 g/L of the target product *Z*11-16OH was obtained. The fatty alcohols were extracted from the yeast biomass using organic solvents and purified on a silica column. The eluted fractions with a high content of the product were pooled and oxidized into corresponding aldehyde using tetrakisacetonitrile copper(I) triflate/TEMPO catalyst system.^20^ The composition of the aldehyde preparation was *Z*11-16Ald, hexadecanal (16Ald), and (*Z*)-hexadec-9-enal (*Z*9-16Ald) in ratio 82:13:5 (Fig. S3, S4). The *Z*11-16Ald is the major and *Z*9-16Ald is the minor pheromone component in *H. armigera* and *C*. *suppressalis*, where the reported ratios between the two pheromone components in *H. armigera* were from 99:1 to 93:7^21–23^, in *C. suppressalis* the reported ratio is 90:10^24^. 16Ald is also present in the pheromone glands of both insect species, but it does not elicit a behavioural response. The biologically produced pheromone composition may thus be close enough and well suited for trapping and mating disruption of these insect species with *Z*11-16Ald as a major and *Z*9-16Ald as a minor pheromone component. The biologically produced pheromone mix was subsequently subjected to activity tests on *H. armigera* in the laboratory and field.

### Electrophysiological responses of male *H. armigera*

We measured the electroantennographic responses of male *H. armigera* adults to the yeast-produced pheromone blend (Bio-Ald), standard compounds, and mixtures of the standards (Fig. 4). Ald mix #1 contained *Z*11-16Ald, *Z*9-16Ald, tetradecanal (14Ald), and pentadecanal (15Ald) (80:5:5:5, respectively). Ald mix #2 contained equal volumes of each of the same components as Ald mix #1 (25:25:25:25 ratio). Bio-Ald elicited the same magnitude of response on the male antenna as Ald mix #1 and significantly higher to that of the equivolume Ald mix #2 and to *Z*9-16Ald, the secondary compound of the *H. armigera* pheromone. The major sex pheromone compound, *Z*11-16Ald, yielded a high antennal response, whereas the minor sex pheromone, *Z*9-16Ald, induced a considerably lower response. The significantly lower response to Ald mix #2 is a clear indication that the antennal response is mainly attributed to *Z*11-16Ald and when its quantity in the mixture is lowered, the antennal response also drops. These results indicate that biologically produced *Z*11-16Ald induces the same magnitude of sensory stimulation as the chemically synthesized *Z*11-16Ald, the major compound of the moth’s native pheromone.

**Fig. 4.**
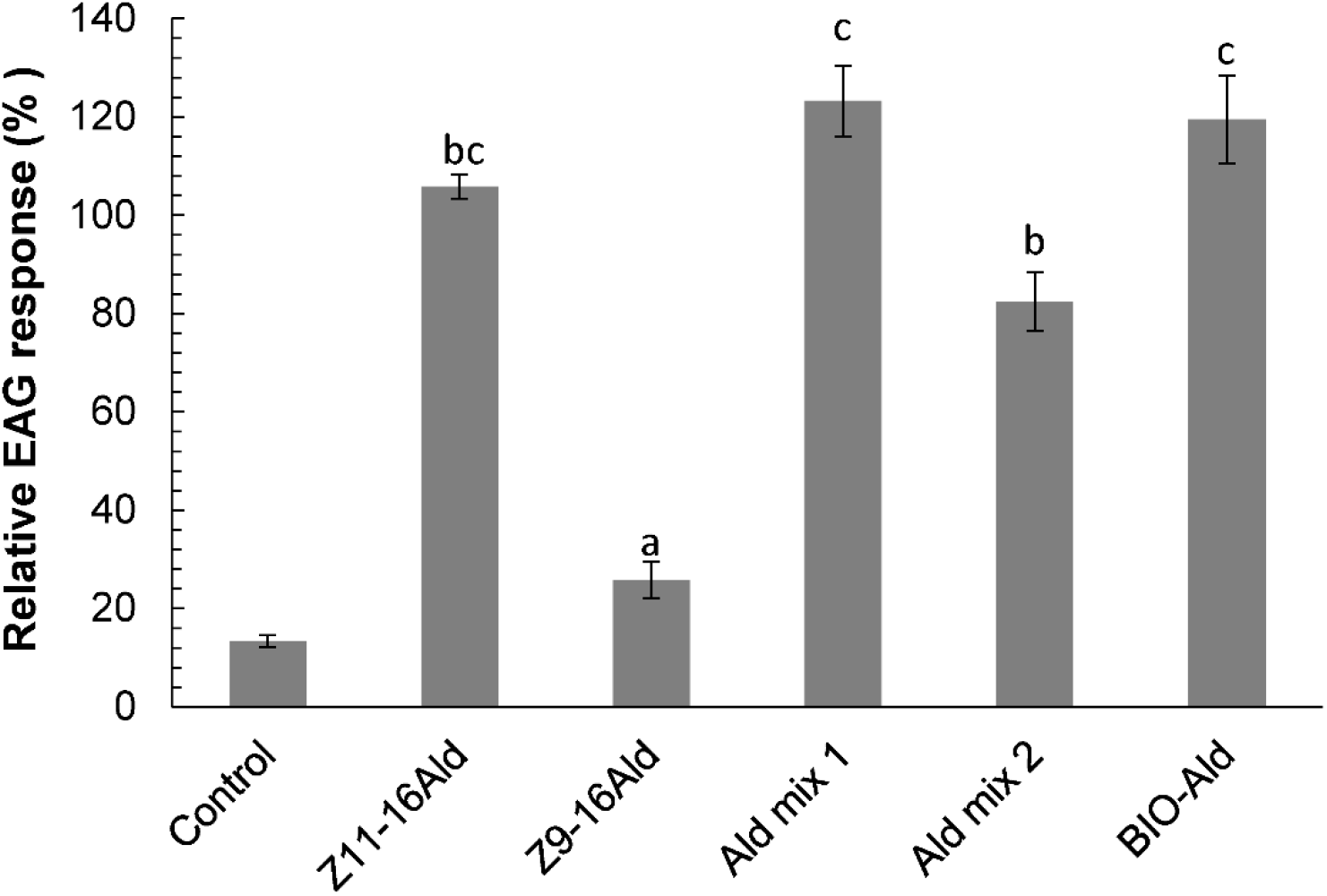
Electrophysiological responses of male *H. armigera* antennae to yeast-produced pheromone blend (Bio-Ald), standard compounds (*Z*11-16Ald, *Z*9-16Ald) and mixtures of the standard compounds (Ald mix#1, Ald mix#2) (±SEM). Ald mix #1 contained *Z*11-16Ald, *Z*9-16Ald, tetradecanal (14Ald), and pentadecanal (15Ald) (80:5:5:5, respectively). Ald mix #2 contained equal volumes of each of the same components as Ald mix #1 (25:25:25:25 ratio). Means followed by the same letter are not significantly different (F=21.491, df=110, P=0.000).

### Monitoring of *H. armigera* flight in the field

Mean weekly male catches in traps baited with yeast-produced pheromone (Bio-Ald) and synthetic pheromone (*Z*11-16Ald: *Z*9-16Ald at 97:3 ratio, Control) dispensers are shown in Fig. 5 for two independent trials. Capture data from the traps indicated that the flight peak occurred on the 2^nd^ week of July (24.7±2.7 males/trap/week for Bio-Ald traps and 22.3±1.8 for the Control traps at Thermi and 19.3±3.0 males/trap/week for Bio-Ald traps and 17.0±3.5 for the Control traps at Lamia). In both regions, the total number of males trapped with the different lures were not significantly different (Bio-Ald: 80.3±4.3 males/trap, Control: 72.0±4.0 males/trap at Thermi and Bio-Ald: 60.3±4.3 males/trap and Control: 57.0±3.6 males/trap at Lamia). It is apparent that the biologically produced pheromone was equally effective under field conditions as the commercially available chemically synthesized one.

**Fig. 5.**
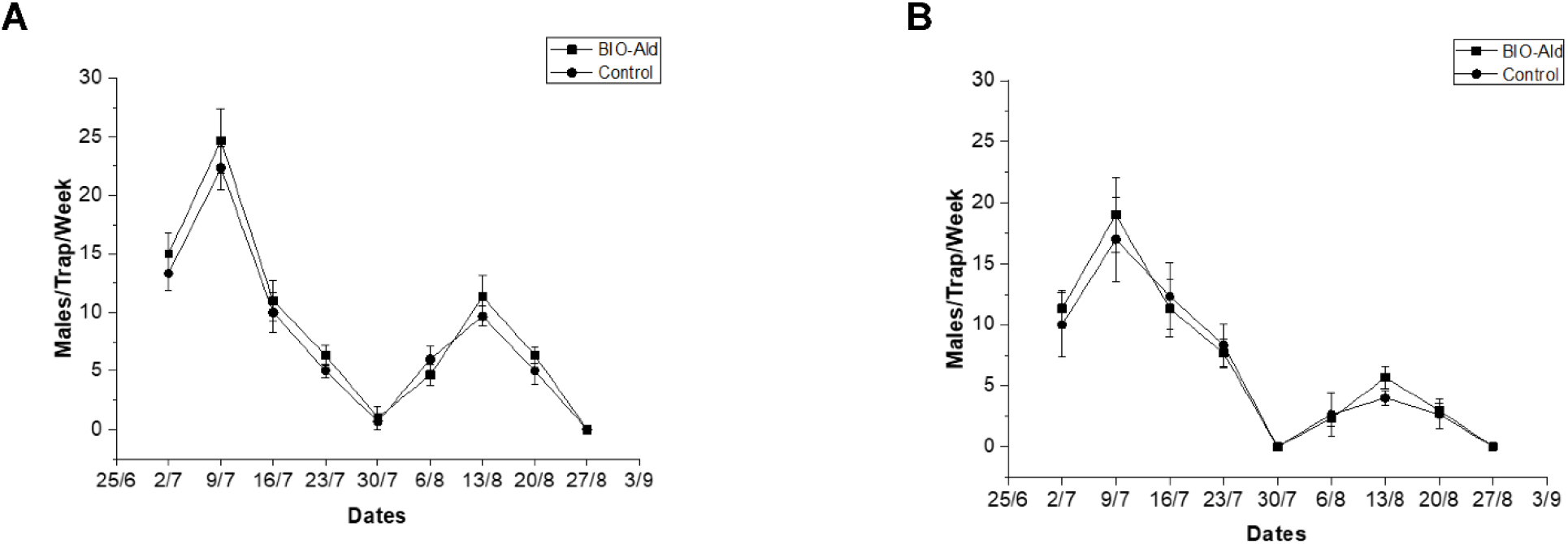
Monitoring of *H. armigera* flight in Greece 2019. Captures of males in pheromone traps baited with commercial synthetic pheromone (Control and yeast-produced pheromone (Bio-Ald) per week (±SEM): (A) in Thermi, Northern Greece (F=1.997, df=5, P=0.231) and (B) Lamia, Central Greece (F=0.189, df=5, P=0.686).

### Conclusions

In summary, we have demonstrated biological production of practically and commercially relevant titres of several lepidopteran sex pheromones in yeast cell factories. A biocatalytic production is particularly advantageous for the production of chemicals for which isomeric composition is critical, such as moth pheromones.^5^ The enzymes can deliver the required stereoisomers, while in chemical synthesis, a mix of isomers is often obtained and may be difficult to separate especially in large quantities. Furthermore, the fermentation is carried out in a cheap medium with glycerol as the sole carbon source, using yeast cells as the only catalyst. This is in contrast to chemical synthesis that will typically require special starting material, expensive catalysts, and several synthesis steps. Reduced production costs and lower environmental impact of the biotech route in comparison to the chemical synthesis have been established for multiple chemicals, particularly for natural products.^25,26^ As an additional advantage, major and minor pheromone components can be produced in a single process in a ratio that is suitable for the target insect. The work creates the foundation for the production of pheromones at a lower cost enabling pheromone-based pest control in row crops, such as rice, cotton, and maize.

## ACKNOWLEDGEMENTS

This project has received funding from the European Union’s Horizon 2020 research and innovation programme under grant agreement No. 760798 (OLEFINE) and from the Bio Based Industries Joint Undertaking (BBI JU) under the European Union’s Horizon 2020 research and innovation programme under grant agreement No: 886662 (PHERA). IB and CL acknowledge the financial support from the Novo Nordisk Foundation under grant agreements No. NNF15OC0016592 and No. NNF10CC1016517 and from Region Sjællands Vækstforum, ViiRS. CL acknowledges funding from the Swedish Foundation for Strategic Research. KP acknowledges the funding from Innovationsfonden under grant agreement number 8053-00179B. LW has received funding from the European Union’s Horizon 2020 research and innovation programme under the Marie Sklodowska-Curie grant agreement No 840491. MK and DR acknowledge Dr. Stefanos Andreadis (Hellenic Agricultural Organization – Demeter) for providing experimental field. The authors thank Volker Zickermann (Goethe-University Frankfurt am Main, Germany), Peter Kötter (Goethe-University Frankfurt am Main), and the Agricultural Research Service (NRRL, USA) for the yeast strains. We thank Dr. Hanne Bjerre Christensen and Dr. Mette Kristensen for the assistance with GC-MS analysis and Dr. Pep Charusanti for help with preparation of pheromone sample for activity tests.

## AUTHOR CONTRIBUTIONS

IB and CL conceived the study. CH, BJD, HLW, MID, KP, KRK, LW, CS and BL performed the experiments on pheromone production in yeasts. LF, MP, and BF carried out fermentation in controlled bioreactors and extracted the pheromones. CB, WU, and AM-N purified and oxidized the sample for activity tests. EK, DR, and MK performed laboratory and field tests of pheromones. IB drafted the manuscript and all the authors have contributed to writing.

## ADDITIONAL INFORMATION

Supplementary Information is available for this paper.

## REFERENCES

1. Reddy, G. V. P. & Guerrero, A. New pheromones and insect control strategies. in Vitamins & Hormones (ed. Litwack, G.) vol. 83 493–519 (Academic Press, 2010).

2. Benelli, G., Lucchi, A., Thomson, D. & Ioriatti, C. Sex pheromone aerosol devices for mating disruption: challenges for a brighter future. Insects 10, 308 (2019).

3. Ioriatti, C. & Lucchi, A. Semiochemical strategies for tortricid moth control in apple orchards and vineyards in Italy. J. Chem. Ecol. 42, 571–583 (2016).

4. Ando, T., Inomata, S. & Yamamoto, M. Lepidopteran sex pheromones. in The chemistry of pheromones and other semiochemicals I (ed. Schulz, S.) 51–96 (Springer, 2004). doi:10.1007/b95449.

5. Löfstedt, C., Wahlberg, N. & Millar, J. G. Evolutionary patterns of pheromone diversity in Lepidoptera. in Pheromone communication in moths: evolution, behavior and application (eds. Allison, J. D. & Carde, R. T.) 43–78 (University of California Press, 2016).

6. Bjostad, L. & Roelofs, W. Sex pheromone biosynthesis in *Trichoplusia ni* - key steps involve delta-11 desaturation and chain-shortening. Science 220, 1387–1389 (1983).

7. El-Sayed, A. The pherobase: database of pheromones and semiochemicals. 2014 http://www.pherobase.com.

8. Hagström, Å. K. et al. A moth pheromone brewery: production of (Z)-11-hexadecenol by heterologous co-expression of two biosynthetic genes from a noctuid moth in a yeast cell factory. Microb. Cell Factories 12, 125 (2013).

9. Jensen, N. B. et al. EasyClone: method for iterative chromosomal integration of multiple genes in *Saccharomyces cerevisiae*. FEMS Yeast Res. 14, 238–248 (2014).

10. Batista-Pereira, L. G., Stein, K., De Paula, A. F., Moreira, J. A., Cruz, I., Figueiredo, M. L. C., Perri, J., Jr., and Corrêa, A. G. Isolation, identification, synthesis, and field evaluation of the sex pheromone of the Brazilian population of *Spodoptera frugiperda*. J. Chem. Ecol. 32, 1085–1099 (2006).

11. Binning, R. R., Coats, J., Kong, X. & Hellmich, R. L. Susceptibility and aversion of *Spodoptera frugiperda* (Lepidoptera: Noctuidae) to Cry1F Bt maize and considerations for insect resistance management. J. Econ. Entomol. 107, 368–374 (2014).

12. Ding, B.-J. et al. The yeast ATF1 acetyltransferase efficiently acetylates insect pheromone alcohols: implications for the biological production of moth pheromones. Lipids 51, 469–475 (2016).

13. Xue, Z. et al. Production of omega-3 eicosapentaenoic acid by metabolic engineering of *Yarrowia lipolytica*. Nat. Biotechnol. 31, 734–740 (2013).

14. Shaw, A. J. et al. Metabolic engineering of microbial competitive advantage for industrial fermentation processes. Science 353, 583–586 (2016).

15. Holkenbrink, C. et al. EasyCloneYALI: CRISPR/Cas9-based synthetic toolbox for engineering of the yeast *Yarrowia lipolytica*. Biotechnol. J. 0, 1700543.

16. Stovicek, V., Borodina, I. & Forster, J. CRISPR–Cas system enables fast and simple genome editing of industrial *Saccharomyces cerevisiae* strains. Metab. Eng. Commun. 2, 13–22 (2015).

17. Iwama, R., Kobayashi, S., Ohta, A., Horiuchi, H. & Fukuda, R. Fatty aldehyde dehydrogenase multigene family involved in the assimilation of n-alkanes in *Yarrowia lipolytica*. J. Biol. Chem. 289, 33275–33286 (2014).

18. Iwama, R., Kobayashi, S., Ohta, A., Horiuchi, H. & Fukuda, R. Alcohol dehydrogenases and an alcohol oxidase involved in the assimilation of exogenous fatty alcohols in *Yarrowia lipolytica*. FEMS Yeast Res. 15, fov014 (2015).

19. Rigouin, C. et al. Production of medium chain fatty acids by *Yarrowia lipolytica*: combining molecular design and TALEN to engineer the fatty acid synthase. ACS Synth. Biol. 6, 1870–1879 (2017).

20. Hoover, J. M. & Stahl, S. S. A highly practical copper(I)/TEMPO catalyst system for chemoselective aerobic oxidation of primary alcohols. J. Am. Chem. Soc. 133, 16901–16910 (2011).

21. Nesbitt, B. F., Beevor, P. S., Hall, D. R. & Lester, R. (Z)-9-Hexadecenal: a minor component of the female sex pheromone of *Heliothis armigera* (Hübner) (Lepidoptera, Noctuidae). Entomol. Exp. Appl. 27, 306–308 (1980).

22. Dunkelblum, E., Gothilf, S. & Kehat, M. Identification of the sex pheromone of the cotton bollworm, *Heliothis armigera*, in Israel. Phytoparasitica 8, 209–211 (1980).

23. Zhang, J.-P., Salcedo, C., Fang, Y.-L., Zhang, R.-J. & Zhang, Z.-N. An overlooked component: (Z)-9-tetradecenal as a sex pheromone in *Helicoverpa armigera*. J. Insect Physiol. 58, 1209–1216 (2012).

24. Tatsuki, S. et al. Sex pheromone of the rice stem borer, *Chilo suppressalis* (WALKER) (Lepidoptera: Pyralidae): the third component, Z-9-hexadecenal. Appl. Entomol. Zool. 18, 443–446 (1983).

25. Pellis, A., Cantone, S., Ebert, C. & Gardossi, L. Evolving biocatalysis to meet bioeconomy challenges and opportunities. New Biotechnol. 40, 154–169 (2018).

26. Jullesson, D., David, F., Pfleger, B. & Nielsen, J. Impact of synthetic biology and metabolic engineering on industrial production of fine chemicals. Biotechnol. Adv. 33, 1395–1402 (2015).

