## Supplemental Information for "Production of moth sex pheromones for pest control by yeast fermentation"

**This PDF file includes:**

Methods
DNA sequences of synthetic genes and plasmids
Figures S1 to S4
Tables S1 to S6
SI References

### Methods

**DNA assembly and yeast strain construction.** Heterologous genes were codon-optimized for *S. cerevisiae* or *Y. lipolytica* and synthesized (GeneArt, Thermo Scientific). The vectors for gene expression and knock-outs were assembled and transformed into yeast according to the published methodologies (1–4). The sequences of the genes and primers, the schemes for gene amplification, cloning and strain assembly are provided in Tables S1-5. Yeast strain *Y. lipolytica* GB20 was a kind gift of Volker Zickermann (Goethe-University Frankfurt am Main, Germany). Yeast strain *S. cerevisiae* CEN.PK102-5B was obtained from Peter Kötter (Goethe-University Frankfurt am Main, Germany). Strain *Y. lipolytica* Y-17536 was received from the Agricultural Research Service (NRRL, USA). For removal of selection markers from the genome of *Y. lipolytica*, we used CreA recombinase gene obtained from plasmid pSH66 (EUROSCARF selection).

**Chemicals and media.** All chemicals were purchased from Sigma-Aldrich. Pheromone standards were purchased from Pherobank. Nourseothricin was from WERNER BioAgents.

**Cultivation of yeast strains.**
Fig. 1b: One individual clone of each strain was inoculated into 2 mL medium (100 g/L glucose, 2 g/L yeast extract, 0.33 g/L $(NH_4)_2SO_4$, 1.33 g/L $MgSO_4 \cdot 7H_2O$ 1.33, 0.267 g/L NaCl , 2 mL/L trace metals solution (4.5 g/L $CaCl_2 \cdot 2H_2O$, 4.5 g/L $ZnSO_4 \cdot 7H_2O$, 3 g/L $FeSO_4 \cdot 7H_2O$, 1 g/L $H_3BO_3$, 1 g/L $MnCl_2 \cdot 2H_2O$, 0.4 g/L $Na_2MoO_4 \cdot 2H_2O$, 0.3 g/L $CoCl_2 \cdot 6H_2O$, 0.1 g/L CuSO4 · $5H_2O$, 0.1 g/L KI, 15 g/L EDTA), 8 mg/L thiamine, 0.67 mg/L biotin, 20 mg/L uracil, 380 mg/L leucine, 76 mg/L histidine, 100 mM potassium hydrogen phthalate buffer) in a 12-mL glass tube (Duran, Wertheim, Germany) with metal labocap lids (Lüdiswiss, Flawil, Switzerland) and incubated for 48 hours at 30°C with shaking at 250 rpm.
Fig 2b: One individual clone of each strain was inoculated into 5 mL YPD medium with 8% glucose (10 g/L yeast extract, 20 g/L peptone, 80 g/L dextrose) in 12-mL glass tubes (Duran, Wertheim, Germany) with metal labocap lids (Lüdiswiss, Flawil, Switzerland) and incubated overnight at 30°C with shaking at 250 rpm. The following day the overnight culture was centrifuged, the supernatant was discarded and the pellet was resuspended in 2 mL nitrogen-limited medium (2.9 g/L $(NH_4)_2SO_4$, 1.7 g/L YNB (without amino acids and ammonium sulphate), 380 mg/L leucine, 76 mg/L lysine, 20 mg/L uracil and 60 g/L glucose). The cultures were incubated for 48 hours at 30°C with shaking at 250 rpm.
Fig. 1d: Three individual colonies of strains expressing desaturases were inoculated into 1 mL selective medium (SC-Ura-Leu) and incubated at 30°C and 300 rpm for 48 h. The cultures were diluted to an OD600 of 0.4 in 5 mL selective medium (SC-Ura-Leu) supplemented with 2 mM $CuSO_4$ and the 0.5 mM methyl tetradecanoate (14Me) (Larodan Fine Chemicals, Sweden). The methyl tetradecanoate stock solution was prepared to a concentration of 100 mM in 96% ethanol. The yeast cultures were incubated at 30°C at 300 rpm for 48 hours.
Fig. 1e: Strains ST4854 and ST5290 were inoculated into 5 mL synthetic complete medium (SC-His-Leu-Trp supplemented with 20 mg/L uracil and 76 mg/L histidine) and cultivated in 12-mL glass tubes (Duran, Wertheim, Germany) with metal labocap lids (Ludiswiss, Flawil, Switzerland) overnight at 30°C with shaking at 250 rpm. The following day the overnight culture was centrifuged, the supernatant was discarded and the pellet was resuspended in 2 mL of mineral medium, which had the composition as described elsewhere[5]. The medium was supplemented with 20 mg/L uracil and 76 mg/L histidine. The cultures were incubated at 30°C with shaking at 250 rpm for 48 hours.
Fig. 2c: One individual clone of each strain was inoculated into 3 mL mineral medium (14.5 g/L $KH_2PO_4$, 0.5 g/L $MgSO_4$, 2 mL/L trace metals solution (4.5 g/L $CaCl_2 \cdot 2H_2O$, 4.5 g/L $ZnSO_4 \cdot 7H_2O$, 3 g/L $FeSO_4 \cdot 7H_2O$, 1 g/L $H_3BO_3$, 1 g/L $MnCl_2 \cdot 2H_2O$, 0.4 g/L $Na_2MoO_4 \cdot 2H_2O$, 0.3 g/L $CoCl_2 \cdot 6H_2O$, 0.1 g/L $CuSO_4 \cdot 5H_2O$, 0.1 g/L KI, 15 g/L EDTA), 1 mL/L vitamin solution (50 mg/L biotin, 200 mg/L p-aminobenzoic acid, 1 g/L nicotinic acid, 1 g/L capantotenate, 1 g/L pyridoxine HCl, 1 g/L thiamine HCl, 25 g/L myo-inositol), 6.9 g/L urea, 50 g/L glycerol, 1.9 g/L leucine, 0.38 g/L uracil, 0.38 g/L lysine) in a 24 deep-well plate with air-penetrable lid (EnzyScreen) and cultivated for 24 hours at 30°C with shaking at 250 rpm The following day 3 mL of mineral medium were inoculated with the preculture in a 24 deep-well plate with air-penetrable lid (EnzyScreen) and cultivated for 48 hours

at 30°C with shaking at 250 rpm. After 24 hours of cultivation, 119 µL of glycerol (corresponding to 50 g/L) were added to each well.

**Metabolite extraction and analysis on GC/MS.**

Fig. 1b, Fig. 1e, Fig. 2b: For extraction, 1 mL of culture was transferred into a 4-mL glass vial and 10 µL of internal standard stock (1 µg/µL methyl (Z)-heptadec-10-enoate in 100% ethanol) was added. The samples were freeze-dried in a freeze dry system (Freezone6 and Stoppering tray dryer, Labconco, Kansas City, USA) at -40°C and 1 mL chloroform:methanol (2:1, v/v) mixture was added to the cell residues. The mixture was vortexed for 45 s and left at room temperature for 4 hours. The organic solvents were slowly evaporated to dryness under a nitrogen stream. One mL of hexane was added to recover the alcohol and acetate content, the samples were vortexed for 10 s, centrifuged and 200 µL of the organic supernatant was transferred to a new glass vial. GC/MS analyses were performed on a Hewlett Packard 6890 GC coupled to HP 5973 mass spectrometer detector. The GC was equipped with an INNOWax column (30 m×0.25 mm×0.25 µm), and helium was used as carrier gas (average velocity: 33 cm/s). The MS was operated in electron impact mode (70eV), scanning between *m/z* 30 and 400, and the injector was configured in splitless mode at 220°C. The oven temperature was set to 80°C for 1 min, then increased at a rate of 10°C /min to 210°C, followed by a hold at 210°C for 15 min, and then increased at a rate of 10°C/min to 230°C followed by a hold at 230°C for 20 min. Compounds were identified by comparison of their retention times and mass spectra with those of the corresponding commercially available standards. Data were analyzed by the Agilent ChemStation software and iWork Numbers. The concentrations of fatty alcohols were calculated using internal standards.

Fig. 1d: One mL of culture was sampled and 3.12 µg of methyl nonadecanoate (19Me) was added as internal standard. Total lipids were extracted using 3.75 mL of methanol/chloroform (2:1, v/v), in a glass vial. One mL of acetic acid (0.15 M) and 1.25 mL of water were added to the tube. Tubes were vortexed vigorously and centrifuged at 2,000 x g for 2 min. The bottom chloroform phase, about 1 mL, containing the total lipids, was transferred to a new glass vial and the solvent was evaporated to dryness. Fatty acid methyl esters (FAMEs) were made from this total lipid extract by acid methanolysis, as follows. One mL of 2% sulfuric acid in methanol (v/v) was added to the tube, vortexed vigorously, and incubated at 90°C for 1 h. After incubation, 1 mL of water was added and mixed well, and then 1 mL of hexane was used to extract the FAMEs. The resulting methyl ester samples were subjected to GC/MS analyses on a Hewlett Packard 6890 GC coupled to a HP 5973 mass selective detector as described above. The monounsaturated fatty acid products were identified by comparing their retention times and mass spectra with those of synthetic standards. Data were analyzed by the ChemStation software (Agilent, Technologies, USA).

**Fatty alcohol degradation analysis.** Cells were cultivated according to the same method as described above for Fig. 2b with the exception that the preculture was incubated for 36 hours (instead of overnight) and *Z*9-14OH and *Z*11-16OH were added to a final concentration of 1 g/L (indicated by +Alc). The concentration of fatty alcohols in the whole broth was determined after 48 hours of incubation in the nitrogen-limited medium. Extraction of samples was performed as following: 100 µL of broth was extracted with 1 mL of ethyl acetate:ethanol (85:15) and using 10 µL of methyl nonadecanoate (19Me, 2 mg/mL) as internal standard. The samples were vortexed for 20 s and incubated for 1 h at room temperature, followed by 5 min of vortexing. 300 µL of $H_2O$ was added to each sample. The samples were vortexed and centrifuged for 5 min at 21°C and 3,000 x *g*. The upper organic phase was analysed via gas chromatography-mass spectrometry (GC/MS). GC/MS analyses were performed on an Agilent 7820A GC coupled to 5977B mass selective detector. The GC was equipped with a split/spitless injector and a DB-Fatwax UI column (30 m×0.25 mm×0.25 µm). The operation parameters were: 1 µL split injection (30:1), injector temperature 220°C and constant flow 1 mL/min helium. The oven temperature was set to 80°C for 1 min, then increased at a rate of 15°C/min to 210°C, followed by a hold at 210°C for 7 min, and then increased at a rate of 20°C/min to 230°C. Fatty alcohols were analysed in selected ion monitoring (SIM) mode using the following mass-to charge-ratios for quantification: 55.1 and 74.1. Compounds were identified by comparison of retention times with those of the corresponding commercially available standards. Data were analysed by the Mass Hunter software B.08.00.

**Analysis of GPAT expression by qRT-PCR.** For qRT-PCR analysis, yeast strains ST5789 (control) and ST5791 (a strain with truncated *GPAT* promoter) were cultivated in triplicates according to method described above for Fig. 2b. The harvesting and pre-treatment before RNA extraction was as described in Dahlin *et al*. (5). Total RNA was isolated using RNeasy Mini Kit (Qiagen) according to manufacturer's instructions. First strand cDNA synthesis was performed using SuperScript™ II Reverse Transcriptase (ThermoFisher Scientific). 20 ng of cDNA was used for qRT-PCR analysis which was done using DyNAmo Flash SYBR Green qPCR Kit on a Stratagene Mx3005P (Agilent Technologies). Relative expression level was calculated using double delta method (ΔΔCt), where ΔΔCt = (ΔCtE − ΔCtC).

**Analysis of lipid content in strain with GPAT downregulation**
Cells cultivated in triplicates according to method described above for Fig. 2b were subjected to FAMEs analysis. 1 mL of broth was transferred to 4 mL glass vials and centrifuged at 3000 g for 5 minutes at room temperature, the supernatant discarded and the cell pellet treated with 1 mL of 1M HCl in methanol, vortexed and incubated at 80 °C for 2 hours. After methanolysis, the mixture was neutralized with 1 mL of 1M NaOH in methanol, then 0.5 mL of saturated NaCl in water was added and 1 mL of hexane was added together with internal standard (19Me). Mixture was vortexed and centrifuged at 3000 g for 5 minutes at room temperature and then the upper phase recovered for GC analysis. GC/MS analysis was done on Agilent 7820A GC system coupled to 5977B mass selective detector. The GC was equipped with DB-Fatwax UI column (30 m×0.25 mm×0.25 µm) and helium was used as carrier gas (average velocity 36.966 cm/sec). The MS was scanning between *m/z* 30 and 350 and the injector was configured to split mode (split ratio 10:1) at 220°C. The oven temperature was set to 80°C for 1 min, then increased at a rate of 15 °C/min to 210°C, followed by a hold at 210°C for 7 min. Then temperature increased at a rate of 20 °C/min to 230°C. For injection 1 µl of sample was used. Compounds were identified by comparison of retention times with those of reference compounds and quantified based on internal standard. For dry cell weight (DW) measurements 1 mL of culture was taken, centrifuged for 5 minutes at 16000 g, supernatant discarded, pellet washed with 1 mL sterile water, centrifuged once again and water discarded. Washed pellets were kept in an oven at 65°C for 48 hours and biomass weighted. Effect of GPAT downregulation was evaluated based on ratio between FAMEs and DW normalized to ST-5789. FAMEs which were included in analysis can be seen in Fig. S2.

**Preparation of yeast-derived biological pheromone sample.** The fermentations were carried out in a BioFlo 415 bioreactor (Eppendorf/NewBrunswick Scientific), equipped with a *in-situ* sterilized 14 L stainless steel vessel (10 L max working volume). pH was controlled at 5.0±0.1 with automated addition of a 1M solution of $H_2SO_4$, and a 4M solution of NaOH. Dissolved oxygen was measured using a polarographic electrode and automatically controlled at 20% saturation by changing the stirring speed of three 6-blade Rushton turbines. Strain *Yarrowia lipolytica* ST6379 was inoculated into 6 L of fermentation medium (2 g/L yeast extract, 13.4 g/L yeast nitrogen base, 0.76 g/L lysine, 0.76 g/L uracil, 0.024 mg/L thiamine, 0.002 g/L biotin, and 50 g/L glycerol). After 20 hours of fermentation, the culture was supplemented with 750 mL nutrient-rich feed (composed of 16.2 g/L yeast extract, 108.6 g/L yeast nitrogen base, 6.2 g/L lysine, 0.2 mg/L thiamine, 0.02 g/L biotin, and 326 g/L glycerol), followed by a pulse of glycerol to a concentration of 50 g/L in the reactor at 32 hours. From 36 hours, glycerol was fed continuously keeping a steady glycerol concentration of 20-30 g/L in the bioreactor. The fermentation lasted a total of 48 hours. The concentration of *Z*11-16:OH was 2.57 g/L.
Liquid-liquid extraction with ethyl acetate was performed on a total of 4.2 L fermentation broth. Fermentation broth was centrifuged at 4000xg for 5 minutes. The supernatant was discarded and the remaining pellet was freeze dried and pulverized. 1 L ethyl acetate was added to the pulverized powder and incubated on a multi-vortexer for 8 h. After filtering off the solvent, the biomass cakes were re-extracted with 0.5 L fresh ethyl acetate. All extracts were combined and the solvent was evaporated to dryness. The extract was resuspended in 25 mL ethyl acetate.
For purification, the crude extract (4.7 g) was passed through a plug of silica gel (approximately 100 g), in a filtration funnel. The silica was washed with hexanes, and then subsequently with a gradient of hexanes:ethyl acetate at the proportions of 95:5 (%) to 80:20 (%), with 5% increments. Fractions were analyzed by TLC eluted with a mixture of hexanes:ethyl acetate in the proportion

80:20 and gas chromatography. The purest fractions were combined and the solvent was evaporated, initially by rotary evaporator and then by means of high vacuum pump. The total amount of purified material recovered was 1.8 g. A fraction of the purified material was transformed into the aldehyde, according to the following protocol: TEMPO (2,2,6,6-Tetramethyl-1-piperidinyloxy, 26 mg) and 1-methylimidazole (28 mg) were added to a well-stirred suspension of the alcohol (800 mg), acetonitrile (3 mL), 2,2′-bipyridyl (26 mg) and tetrakisacetonitrile copper(I) triflate (62 mg). The mixture was stirred at room temperature and open atmosphere for 2 hours and the completion of the reaction was verified by gas chromatography. The solvent was evaporated and the recovered material was extracted with hexanes, water and $NaHCO_3$ aqueous solution. The organic phase was dried with $MgSO_4$, filtered and the solvent was evaporated under reduced pressure. The resulting material (0.58 g) was dissolved in 10 mL ethyl acetate and analyzed as follows. Three replicates 200 µl of the solution were transferred into 5-mL volumetric flasks, and ethyl acetate was added to complete the volume to the meniscus line. The contents of the flasks were mixed by swirling or inverting the flasks up and down several times and aliquots of 1.5 mL were transferred to autosampler vials for GC injection. Analysis was performed in a Agilent 7890 equipped with an FID detector and an HP-5 capillary column. The oven temperature program involved an initial temperature of 115°C, increased at a rate of 40°C/min to 162°C and held for 3 minutes. The temperature was finally increased to 40°C/min to 280°C and held for 3 minutes. Three concentrations of technical grade *Z*11-16Ald in ethyl acetate were prepared and the solutions were also injected into the GC to create a calibration curve and the equation of the line was used for quantitation of *Z*11-16Ald, according to Table S6. The concentration of Z11-16Ald was determined to be 35.33 mg/mL based on three replicates with the RSD of 0.72% between replicates.

**Electrophysiological responses of male *H. armigera*.** The antennal responses of *Helicoverpa armigera* male adults to the pheromone blend produced from yeast fermentation were evaluated by electroantennography (EAG) using a commercially available electroantennographic system (Syntech, The Netherlands). Antennae of a virgin, two-to-three-days-old male adult were used. The signal was amplified and detected with a data acquisition controller (IDAC-4, Syntech, The Netherlands).

Yeast-produced pheromone blend (Bio-Ald), standard compounds and mixtures of the standards were tested at a total of 39 antennal preparations. As standard compounds, the two pheromone components of *H. armigera* pheromone, *Z*11-16:Ald and *Z*9-16:Ald, were used. Two mixtures of the two monounsaturated aldehydes with tetradecanal and pentadecanal were also tested on grounds that presence of tetradecanal and pentadecanal has been verified in the pheromone blend produced by yeast fermentation (MS identification, data not shown). The two blends tested were Ald mix 1: *Z*11-16:Ald, *Z*9-16:Ald, 14:Ald, 15:Ald at 80:5:5:5 ratio and Ald mix 2: *Z*11-16:Ald, *Z*9-16:Ald, 14:Ald, 15:Ald at 1:1:1:1 ratio. Ald mix 1 blend constituents approximate the abundances found in the yeast-produced pheromone blend and notably the ratio of the two monounsaturated aldehydes (94:6) approximate the optimal pheromone blend ratio for *H. armigera.*

Aliquots of 1 µg of each of the compounds (or mixtures) was presented to the male antenna. Stimuli were provided as 0.3 s air puffs into a continuous flow of filtered and humidified air. The air flow, at 25 $cm^3$/s rate, was generated by an air stimulus controller (CS-55, Syntech, The Netherlands). At least 1 min was allowed between successive stimulations in order to allow the antenna to recover. Control stimulus consisted of filter paper and solvent (*n*-pentane). A reference stimulus, consisting of a 1 µg of *Z*11-16:Ald (the major sex pheromone of *H. armigera*), was provided at regular intervals during each recording session. The EAG response to each reference stimulus was defined as 100%, and all responses to the test stimuli between adjacent references were normalized in % relative to the references.

**Monitoring of *H. armigera* flight in the field.** Field trials were conducted in Thermi (northern Greece 40°32'11.6"N 23°00'08.0"E) and in Lamia (central Greece 38°87'64.1"N 22°36'81.3"E) on experimental pesticide-free cotton fields (planted area 1.5 ha each). The two discrete geographical regions having similar meteorological conditions (temperature 27-28ºC; rainfall 2-0.5 mm for July and August respectively).

Dispensers used as control were grey rubber septa (bromobutyl elastomers) loaded with 2mg of *H. armigera* pheromone blend *Z*11-16:Ald : *Z*9-16:Ald at 97:3 ratio) (provided by Novagrica Hellas

SA). Treatment dispensers (Bio-Ald) were similarly bromobutyl elastomers loaded with 2 mg of the yeast-produced pheromone. BHT and bumetrizole were added as antioxidant and UV absorber respectively at 5% w/w. Six funnel traps (three dispensers loaded with commercially available pheromone, control, and three loaded with the yeast-produced pheromone (Bio-Ald) were installed at 1.2 m height. Baited traps were in operation from early July until early September, the traps were rotated clockwise weekly, and males captured were recorded once per week and removed. Pheromone dispensers were renewed every month (6).

**Statistical Analyses.** The electrophysiological and field data were subjected to analysis of variance (ANOVA) (SAS Institute, 2000). The means of electrophysiological data were separated using the Tukey (honestly significant difference, HSD) test at P = 0.05. The field data presented as means of male catches per trap per week.

**DNA sequences of synthetic genes and plasmids**

>SEQ ID NO: 1. Fatty acyl-CoA reductase from *Agrotis segetum*
ATGCCAGTCTTGACTTCTAGAGAAGACGAAAAATTGTCCGTCCCAGAATTTTACGCTGGTAA
GTCTATTTTTGTTACCGGTGGTACTGGTTTCTTGGGTAAGGTTTTTATCGAAAAGTTGTTGTA
CTGCTGCCCAGATATCGATAAGATCTACATGTTGATCAGAGAAAAAAAGAACTTGTCCATCG
ACGAAAGAATGTCCAAGTTTTTGGATGACCCTTTGTTCTCCAGATTGAAAGAAGAAAGACCA
GGTGACTTGGAAAAGATCGTTTTGATTCCAGGTGATATTACCGCTCCTAATTTGGGTTTGTCT
GCTGAAAACGAAAGAATCTTGTTGGAAAAGGTCAGTGTCATTATTAACTCTGCTGCTACCGTT
AAGTTCAACGAACCATTGCCAATTGCTTGGAAGATTAACGTTGAAGGTACTAGAATGTTGTTG
GCCTTGTCTAGAAGAATGAAGAGAATCGAAGTTTTCATCCATATCTCCACCGCTTACTCTAAT
GCTTCTTCTGATAGAATTGTCGTTGACGAAATCTTGTATCCAGCTCCAGCTGATATGGATCAA
GTTTATCAATTGGTTAAGGACGGTGTCACTGAAGAAGAAACCGAAAGATTATTGAACGGTTT
GCCAAACACTTACACTTTCACTAAGGCTTTGACCGAACATTTGGTTGCTGAACATCAAACTTA
CGTTCCAACCATTATCATCAGACCATCTGTTGTTGCCTCCATTAAGGATGAACCTATTAGAGG
TTGGTTGTGTAATTGGTTTGGTGCTACTGGTATTTCTGTTTTCACTGCTAAGGGTTTGAACAG
AGTTTTGTTGGGTAAAGCCTCTAACATCGTTGATGTTATCCCAGTTGATTACGTTGCCAACTT
GGTTATAGTTGCTGGTGCTAAATCTGGTGGTCAAAAGTCTGATGAATTGAAAATCTACAACTG
CTGCTCCTCTGACTGTAATCCAGTTACTTTGAAGAAGATCATCAAAGAATTCACCGAAGATAC
CATCAAGAACAAGTCCCATATTATGCCATTGCCAGGTTGGTTCGTTTTTACTAAGTACAAATG
GTTGTTGACTTTGTTGACCATCATCTTCCAAATGTTGCCAATGTATTTGGCCGATGTTTACAG
AGTCTTGACCGGTAAAATTCCAAGATATATGAAGTTGCACCACTTGGTCATTCAAACCAGATT
GGGTATTGATTTCTTCACCTCTCATTCTTGGGTTATGAAGACCGATAGAGTCAGAGAATTATT
CGGTTCTTTGTCCTTGGCCGAAAAACACATGTTTCCATGTGATCCATCTTCCATTGATTGGAC
CGATTACTTGCAATCTTACTGCTATGGTGTCAGAAGATTCTTAGAAAAGAAGAAGTAA

>SEQ ID NO: 2. Fatty acyl-CoA reductase from *Heliothis subflexa*
ATGGTTGTCTTGACCTCCAAAGAAACTAAGCCATCTGTTGCTGAATTTTACGCTGGTAAGTCT
GTTTTCATTACTGGTGGTACTGGTTTCTTGGGTAAGGTTTTCATTGAAAAGTTGTTGTACTCC
TGCCCAGATATCGGTAATATCTACATGTTGATCAGAGAAAAGAAGGGTTTGTCCGTTTCCGA
AAGAATCAAGCACTTTTTGGATGATCCTTTGTTCACCAGATTGAAAGAAAAAGACCAGCCGA
CTTGGAAAAGATCGTTTTGATTCCAGGTGATATTACTGCTCCAGATTTGGGTATTACCTCCGA
AAACGAAAAGATGTTGATCGAAAAGGTCAGTGTCATTATTCATTCTGCTGCTACCGTTAAGTT
CAACGAACCATTGCCAACTGCTTGGAAGATTAACGTTGAAGGTACTAGAATGATGTTGGCCT
TGTCTAGAAGAATGAAGAGAATCGAAGTTTTCATCCATATCTCTACCGCTTACACTAACACCA
ACAGAGAAGTTGTTGACGAAATCTTGTATCCAGCTCCAGCTGATATTGATCAAGTTCACCAAT
ATGTTAAGGACGGTATCTCTGAAGAAGAAACTGAAAAAATCTTGAACGGTAGACCAAACACT
TACACTTTCACTAAGGCTTTGACCGAACATTTGGTTGCTGAAAATCAAGCTTACGTTCCAACC
ATTATCGTTAGACCATCAGTTGTTGCTGCCATTAAGGATGAACCTATTAAGGGTTGGTTGGGT
AATTGGTATGGTGCTACAGGTTTGACTGTTTTTACTGCTAAGGGTTTGAACAGAGTTATCTAC
GGTCACTCTTCTAACATCGTTGATTTGATCCCAGTTGATTACGTTGCCAACTTGGTTATTGCT
GCTGGTGCTAAATCTTCTAAGTCTACTGAATTGAAGGTCTACAACTGCTGTTCTTCTGCTTGT
AACCCAATTACTATCGGTAAGTTGATGTCCATGTTTGCTGAAGATGCTATCAAGCAAAAGTCT
TACGCTATGCCATTGCCAGGTTGGTACATTTTTACTAAGTACAAGTGGTTGGTCTTGTTGTTG
ACCATTTTGTTCCAAGTTATTCCAGCCTACATTACCGACTTGTACAGACATTTGATTGGTAAG
AACCCAAGATATATCAAGTTGCAATCCTTGGTCAATCAAACCAGATCCTCCATTGATTTCTTC
ACCAACCATTCTTGGGTTATGAAGGCTGATAGAGTCAGAGAATTATTCGCTTCTTTGTCTCCA
GCAGATAAGTACTTGTTTCCATGTGATCCAGTCAACATCAATTGGAGACAATATATCCAAGAT
TACTGCTGGGGTGTTAGACATTTCTTGGAAAAAAAGACTTAA

>SEQ ID NO: 3. Fatty acyl-CoA reductase from *Helicoverpa assulta*
ATGGTTGTCTTGACCTCCAAAGAAACTAAGCCATCTGTTGCTGAATTTTACGCTGGTAAGTCT
GTTTTCATTACTGGTGGTACTGGTTTCTTGGGTAAGATCTTCATTGAAAAGTTGTTGTACTCC
TGCCCAGATATCGGTAATATCTACATGTTGATCAGAGAAAAGAAGGGTTTGTCCGTTTCCGA
AAGAATCAAGCAATTTTTGGATGACCCTTTGTTCACCAGATTGAAAGAAAAAGACCAGCCG
ACTTGGAAAAGATCGTTTTGATTCCAGGTGATATTACTGCTCCAGATTTGGGTATTACCTCCG

AAAACGAAAAGATGTTGATCGAAAAGGTCAGTGTCATTATTCATTCTGCTGCTACCGTTAAGT
TCAACGAACCATTGCCAACTGCTTGGAAGATTAACGTTGAAGGTACTAGAATGATGTTGGCC
TTGTCTAGAAGAATGAAGAGAATCGAAGTTTTCATCCATATCTCTACCGCTTACACTAACACC
AACAGAGAAGTTGTTGACGAAATCTTGTATCCAGCTCCAGCTGATATTGATCAAGTTCACCAA
TATGTTAAGGACGGTATCTCTGAAGAAGAAACTGAAAAAATCTTGAACGGTAGACCAAACAC
TTACACTTTCACTAAGGCTTTGACCGAACATTTGGTTGCTGAAAATCAAGCTTACGTTCCAAC
CATTATCGTTAGACCATCAGTTGTTGCTGCCATTAAGGATGAACCTATTAAGGGTTGGTTGG
GTAATTGGTATGGTGCTACAGGTTTGACTGTTTTTACTGCTAAGGGTTTGAACAGAGTTATCT
ACGGTCATTCCTCTTACATCGTTGATTTGATCCCAGTTGATTACGTTGCCAACTTGGTTATTG
CTGCTGGTGCTAAATCTTCTAAGTCTACTGAATTGAAGGTCTACAACTGCTGTTCTTCTGCTT
GTAACCCAATTACTATCGGTAAGTTGATGTCCATGTTTGCTGAAGATGCTATCAAGCAAAAGT
CTTACGCTATGCCATTGCCAGGTTGGTATGTTTTTACAAAGTACAAGTGGTTGGTCTTGTTGT
TGACCATTTTGTTCCAAGTTATTCCAGCCTACATTACCGACTTGTACAGACATTTGATTGGTA
AGAACCCAAGATATATCAAGTTGCAATCCTTGGTCAATCAAACCAGATCCTCCATTGATTTCT
TCACCTCTCATTCTTGGGTTATGAAGGCTGATAGAGTCAGAGAATTATTCGCTTCTTTGTCTC
CAGCAGATAAGTACTTGTTTCCATGTGATCCAACCGATATTAACTGGACCCATTACATTCAAG
ATTACTGCTGGGGTGTTAGACACTTCTTGGAAAAAAAGACTACCAACAAGTAA

>SEQ ID NO: 4. Δ11-desaturase from *Amyelois transitella*
ATGGTTCCAAACAAGGGTTCCTCTGATGTTTTGTCTGAACATTCTGAACCACAATTCACCAAG
TTGATTGCTCCACAAGCTGGTCCAAGAAAGTACAAAATCGTTTACAGAAACTTGTTGACCTTC
GGTTACTGGCATTTGTCTGCTGTTTATGGTTTGTACTTGTGTTTCACTTGTGCTAAGTGGGCT
ACTATTTTGTTCGCTTTCTTCTTGTACGTTATCGCCGAAATTGGTATTACTGGTGGTGCTCAT
AGATTATGGGCTCATAGAACTTACAAAGCCAAGTTGCCATTGGAAATCTTGTTGTTGATCATG
AACTCCATTGCCTTCCAAGATACTGCTTTTACTTGGGCTAGAGATCATAGATTGCATCACAAG
TACTCTGATACTGATGCTGATCCACATAATGCTACTAGAGGTTTCTTCTACTCTCATGTTGGT
TGGTTGTTGGTTAAGAAACACCCAGAAGTTAAGGCTAGAGGTAAGTACTTGTCTTTGGATGA
CTTGAAGAACAACCCTTTGTTGAAGTTCCAAAAGAAGTACGCCATTTTGGTCATTGGTACTTT
GTGCTTTTTGATGCCAACTTTCGTTCCAGTTTACTTTTGGGGTGAAGGTATTTCTACTGCCTG
GAACATTAACTTGTTAAGATACGTCATGAACTTGAACATGACCTTTTTGGTTAACTCCGCTGC
TCATATTTTTGGTAACAAGCCATACGATAAGTCTATCGCCTCTGTTCAAAACATCTCTGTTTCT
TTGGCTACTTTCGGTGAAGGTTTCCATAACTACCATCATACTTATCCATGGGATTACAGAGCT
GCTGAATTGGGTAACAATAGATTGAATATGACCACCGCCTTCATTGATTTCTTTGCTTGGATT
GGTTGGGCCTACGATTTGAAATCTGTTCCACAAGAAGCTATTGCTAAGAGATGTGCTAAAAC
TGGTGATGGTACTGATATGTGGGGTAGAAAGAGATGA

>SEQ ID NO: 5. Fatty acyl-CoA reductase from *Helicoverpa armigera*
ATGGTTGTCTTGACCTCCAAAGAAACTAAGCCATCTGTTGCTGAATTTTACGCTGGTAAGTCT
GTTTTCATTACTGGTGGTACTGGTTTCTTGGGTAAGGTTTTCATTGAAAAGTTGTTGTACTCC
TGCCCAGATATCGGTAATATCTACATGTTGATCAGAGAAAAGAAGGGTTTGTCCGTTTCCGA
AAGAATCAAGCACTTTTTGGATGATCCTTTGTTCACCAGATTGAAAGAAAAAGACCAGCCGA
CTTGGAAAAGATCGTTTTGATTCCAGGTGATATTACTGCTCCAGATTTGGGTATTACCTCCGA
AAACGAAAAGATGTTGATCGAAAAGGTCAGTGTCATTATTCATTCTGCTGCTACCGTTAAGTT
CAACGAACCATTGCCAACTGCTTGGAAGATTAACGTTGAAGGTACTAGAATGATGTTGGCCT
TGTCTAGAAGAATGAAGAGAATCGAAGTTTTCATCCATATCTCTACCGCTTACACTAACACCA
ACAGAGAAGTTGTTGACGAAATCTTGTATCCAGCTCCAGCTGATATTGATCAAGTTCACAGAT
ATGTTAAGGACGGTATCTCTGAAGAAGAAACTGAAAAAATCTTGAACGGTAGACCAAACACT
TACACTTTCACTAAGGCTTTGACCGAACATTTGGTTGCTGAAAATCAAGCTTACGTTCCAACC
ATTATCGTTAGACCATCAGTTGTTGCTGCCATTAAGGATGAACCTATTAAGGGTTGGTTGGGT
AATTGGTATGGTGCTACAGGTTTGACTGTTTTTACTGCTAAGGGTTTGAACAGAGTTATCTAC
GGTCACTCTTCTAACATCGTTGATTTGATCCCAGTTGATTACGTTGCCAACTTGGTTATTGCT
GCTGGTGCTAAATCTTCTAAGTCTACTGAATTGAAGGTCTACAACTGCTGTTCTTCTGCTTGT
AACCCAATTACTATCGGTAAGTTGATGTCCATGTTTGCTGAAGATGCTATCAAGCAAAAGTCT
TACGCTATGCCATTGCCAGGTTGGTACATTTTTACTAAGTACAAGTGGTTGGTCTTGTTGTTG
ACCATTTTGTTCCAAGTTATTCCAGCCTACATTACCGACTTGTACAGACATTTGATTGGTAAG
AACCCAAGATATATCAAGTTGCAATCCTTGGTCAATCAAACCAGATCCTCCATTGATTTCTTC

ACCTCTCATTCTTGGGTTATGAAGGCTGATAGAGTCAGAGAATTATTCGCTTCTTTGTCTCCA
GCAGATAAGTACTTGTTTCCATGTGATCCAACCGATATTAACTGGACCCATTACATTCAAGAT
TACTGCTGGGGTGTTAGACATTTCTTGGAAAAAAAAAGCTACGAATAA

>SEQ ID NO: 6. Fatty acyl-CoA reductase from *Helicoverpa armigera* with modified C-terminus
ATGGTTGTCTTGACCTCCAAAGAAACTAAGCCATCTGTTGCTGAATTTTACGCTGGTAAGTCT
GTTTTCATTACTGGTGGTACTGGTTTCTTGGGTAAGGTTTTCATTGAAAAGTTGTTGTACTCC
TGCCCAGATATCGGTAATATCTACATGTTGATCAGAGAAAAGAAGGGTTTGTCCGTTTCCGA
AAGAATCAAGCACTTTTTGGATGATCCTTTGTTCACCAGATTGAAAGAAAAAAGACCAGCCGA
CTTGGAAAAGATCGTTTTGATTCCAGGTGATATTACTGCTCCAGATTTGGGTATTACCTCCGA
AAACGAAAAGATGTTGATCGAAAAGGTCAGTGTCATTATTCATTCTGCTGCTACCGTTAAGTT
CAACGAACCATTGCCAACTGCTTGGAAGATTAACGTTGAAGGTACTAGAATGATGTTGGCCT
TGTCTAGAAGAATGAAGAGAATCGAAGTTTTCATCCATATCTCTACCGCTTACACTAACACCA
ACAGAGAAGTTGTTGACGAAATCTTGTATCCAGCTCCAGCTGATATTGATCAAGTTCACAGAT
ATGTTAAGGACGGTATCTCTGAAGAAGAAACTGAAAAAATCTTGAACGGTAGACCAAACACT
TACACTTTCACTAAGGCTTTGACCGAACATTTGGTTGCTGAAAATCAAGCTTACGTTCCAACC
ATTATCGTTAGACCATCAGTTGTTGCTGCCATTAAGGATGAACCTATTAAGGGTTGGTTGGGT
AATTGGTATGGTGCTACAGGTTTGACTGTTTTTACTGCTAAGGGTTTGAACAGAGTTATCTAC
GGTCACTCTTCTAACATCGTTGATTTGATCCCAGTTGATTACGTTGCCAACTTGGTTATTGCT
GCTGGTGCTAAATCTTCTAAGTCTACTGAATTGAAGGTCTACAACTGCTGTTCTTCTGCTTGT
AACCCAATTACTATCGGTAAGTTGATGTCCATGTTTGCTGAAGATGCTATCAAGCAAAAGTCT
TACGCTATGCCATTGCCAGGTTGGTACATTTTTACTAAGTACAAGTGGTTGGTCTTGTTGTTG
ACCATTTTGTTCCAAGTTATTCCAGCCTACATTACCGACTTGTACAGACATTTGATTGGTAAG
AACCCAAGATATATCAAGTTGCAATCCTTGGTCAATCAAACCAGATCCTCCATTGATTTCTTC
ACCTCTCATTCTTGGGTTATGAAGGCTGATAGAGTCAGAGAATTATTCGCTTCTTTGTCTCCA
GCAGATAAGTACTTGTTTCCATGTGATCCAACCGATATTAACTGGACCCATTACATTCAAGAT
TACTGCTGGGGTGTTAGACATTTCTTGGAACATGATGAATTGTAA

>SEQ ID NO: 7. Fatty acyl-CoA reductase from *Heliothis subflexa* with modified C-terminus
ATGGTTGTCTTGACCTCCAAAGAAACTAAGCCATCTGTTGCTGAATTTTACGCTGGTAAGTCT
GTTTTCATTACTGGTGGTACTGGTTTCTTGGGTAAGGTTTTCATTGAAAAGTTGTTGTACTCC
TGCCCAGATATCGGTAATATCTACATGTTGATCAGAGAAAAGAAGGGTTTGTCCGTTTCCGA
AAGAATCAAGCACTTTTTGGATGATCCTTTGTTCACCAGATTGAAAGAAAAAAGACCAGCCGA
CTTGGAAAAGATCGTTTTGATTCCAGGTGATATTACTGCTCCAGATTTGGGTATTACCTCCGA
AAACGAAAAGATGTTGATCGAAAAGGTCAGTGTCATTATTCATTCTGCTGCTACCGTTAAGTT
CAACGAACCATTGCCAACTGCTTGGAAGATTAACGTTGAAGGTACTAGAATGATGTTGGCCT
TGTCTAGAAGAATGAAGAGAATCGAAGTTTTCATCCATATCTCTACCGCTTACACTAACACCA
ACAGAGAAGTTGTTGACGAAATCTTGTATCCAGCTCCAGCTGATATTGATCAAGTTCACCAAT
ATGTTAAGGACGGTATCTCTGAAGAAGAAACTGAAAAAATCTTGAACGGTAGACCAAACACT
TACACTTTCACTAAGGCTTTGACCGAACATTTGGTTGCTGAAAATCAAGCTTACGTTCCAACC
ATTATCGTTAGACCATCAGTTGTTGCTGCCATTAAGGATGAACCTATTAAGGGTTGGTTGGGT
AATTGGTATGGTGCTACAGGTTTGACTGTTTTTACTGCTAAGGGTTTGAACAGAGTTATCTAC
GGTCACTCTTCTAACATCGTTGATTTGATCCCAGTTGATTACGTTGCCAACTTGGTTATTGCT
GCTGGTGCTAAATCTTCTAAGTCTACTGAATTGAAGGTCTACAACTGCTGTTCTTCTGCTTGT
AACCCAATTACTATCGGTAAGTTGATGTCCATGTTTGCTGAAGATGCTATCAAGCAAAAGTCT
TACGCTATGCCATTGCCAGGTTGGTACATTTTTACTAAGTACAAGTGGTTGGTCTTGTTGTTG
ACCATTTTGTTCCAAGTTATTCCAGCCTACATTACCGACTTGTACAGACATTTGATTGGTAAG
AACCCAAGATATATCAAGTTGCAATCCTTGGTCAATCAAACCAGATCCTCCATTGATTTCTTC
ACCAACCATTCTTGGGTTATGAAGGCTGATAGAGTCAGAGAATTATTCGCTTCTTTGTCTCCA
GCAGATAAGTACTTGTTTCCATGTGATCCAGTCAACATCAATTGGAGACAATATATCCAAGAT
TACTGCTGGGGTGTTAGACATTTCTTGCATGATGAATTGTAA

>SEQ ID NO: 8. Fatty acyl-CoA reductase from *Helicoverpa assulta* with modified C-terminus
ATGGTTGTCTTGACCTCCAAAGAAACTAAGCCATCTGTTGCTGAATTTTACGCTGGTAAGTCT
GTTTTCATTACTGGTGGTACTGGTTTCTTGGGTAAGATCTTCATTGAAAAGTTGTTGTACTCC
TGCCCAGATATCGGTAATATCTACATGTTGATCAGAGAAAAGAAGGGTTTGTCCGTTTCCGA

AAGAATCAAGCAATTTTTGGATGACCCTTTGTTCACCAGATTGAAAGAAAAAAGACCAGCCG
ACTTGGAAAAGATCGTTTTGATTCCAGGTGATATTACTGCTCCAGATTTGGGTATTACCTCCG
AAAACGAAAAGATGTTGATCGAAAAGGTCAGTGTCATTATTCATTCTGCTGCTACCGTTAAGT
TCAACGAACCATTGCCAACTGCTTGGAAGATTAACGTTGAAGGTACTAGAATGATGTTGGCC
TTGTCTAGAAGAATGAAGAGAATCGAAGTTTTCATCCATATCTCTACCGCTTACACTAACACC
AACAGAGAAGTTGTTGACGAAATCTTGTATCCAGCTCCAGCTGATATTGATCAAGTTCACCAA
TATGTTAAGGACGGTATCTCTGAAGAAGAAACTGAAAAAATCTTGAACGGTAGACCAAACAC
TTACACTTTCACTAAGGCTTTGACCGAACATTTGGTTGCTGAAAATCAAGCTTACGTTCCAAC
CATTATCGTTAGACCATCAGTTGTTGCTGCCATTAAGGATGAACCTATTAAGGGTTGGTTGG
GTAATTGGTATGGTGCTACAGGTTTGACTGTTTTTACTGCTAAGGGTTTGAACAGAGTTATCT
ACGGTCATTCCTCTTACATCGTTGATTTGATCCCAGTTGATTACGTTGCCAACTTGGTTATTG
CTGCTGGTGCTAAATCTTCTAAGTCTACTGAATTGAAGGTCTACAACTGCTGTTCTTCTGCTT
GTAACCCAATTACTATCGGTAAGTTGATGTCCATGTTTGCTGAAGATGCTATCAAGCAAAAGT
CTTACGCTATGCCATTGCCAGGTTGGTATGTTTTTACAAAGTACAAGTGGTTGGTCTTGTTGT
TGACCATTTTGTTCCAAGTTATTCCAGCCTACATTACCGACTTGTACAGACATTTGATTGGTA
AGAACCCAAGATATATCAAGTTGCAATCCTTGGTCAATCAAACCAGATCCTCCATTGATTTCT
TCACCTCTCATTCTTGGGTTATGAAGGCTGATAGAGTCAGAGAATTATTCGCTTCTTTGTCTC
CAGCAGATAAGTACTTGTTTCCATGTGATCCAACCGATATTAACTGGACCCATTACATTCAAG
ATTACTGCTGGGGTGTTAGACACTTCTTGGAACATGATGAATTGTAA

>SEQ ID NO: 9. Δ11-desaturase from *A. segetum*
ATGGCTCAAGGTGTCCAAACAACTACGATATTGAGGGAGGAAGAGCCGTCATTGACTTTCGT
GGTACCTCAAGAACCGAGAAAGTATCAAATCGTGTACCCAAACCTTATCACATTTGGGTACT
GGCATATAGCTGGTTTATACGGGCTATATTTGTGCTTTACTTCGGCAAAATGGCAAACAATTT
TATTCAGTTTCATGCTCGTTGTGTTAGCAGAGTTGGGAATAACAGCCGGCGCTCACAGGTTA
TGGGCCCACAAAACATATAAAGCGAAGCTTCCCTTACAAATTATCCTGATGATACTGAACTCC
ATTGCCTTCCAAAATTCCGCCATTGATTGGGTGAGGGACCACCGTCTCCATCATAAGTACAG
TGACACTGATGCAGACCCTCACAATGCTACTCGTGGTTTCTTCTATTCTCATGTTGGATGGTT
GCTCGTAAGAAAACATCCAGAAGTCAAGAGACGTGGAAAGGAACTTGACATGTCTGATATTT
ACAACAATCCAGTGCTGAGATTTCAAAAGAAGTATGCTATACCCTTCATCGGGGCAATGTGC
TTCGGATTACCAACTTTTATCCCTGTTTACTTCTGGGGAGAAACCTGGAGTAATGCTTGGCAT
ATCACCATGCTTCGGTACATCCTCAACCTAAACATTACTTTCCTGGTCAACAGTGCTGCTCAT
ATCTGGGGATACAAACCTTATGACATCAAAATATTGCCTGCCCAAAATATAGCAGTTTCCATA
GTAACCGGCGGCGAAGTTTCCATAACTACCACCACGTTTTTTCCTTGGGATTATCGTGCAGC
AGAATTGGGGAACAATTATCTTAATTTGACGACTAAGTTCATAGATTTCTTCGCTTGGATCGG
ATGGGCTTACGATCTTAAGACGGTGTCCAGTGATGTTATAAAAAGTAAGGCGGAAAGAACTG
GTGATGGGACGAATCTTTGGGGTTTAGAAGACAAAGGTGAAGAAGATTTTTTGAAAATCTGG
AAAGACAATTAA

>SEQ ID NO: 10. Δ11-desaturase from *Spodoptera littoralis*
ATGGCTCAATGTGTTCAAACCACCACCATCTTGGAACAAAAAGAAGAAAAGACCGTTACCTT
GTTGGTTCCACAAGCTGGTAAAAGAAAGTTCGAAATCGTCTACTTCAACATCATTACCTTCGC
CTATTGGCATATTGCTGGTTTGTACGGTTTGTATTTGTGTTTCACTTCTACTAAGTGGGCTAC
CGTTTTGTTCTCTTTCTTCTTGTTCGTTGTTGCCGAAGTTGGTGTTACTGCTGGTTCTCATAG
ATTGTGGTCACATAAGACTTACAAGGCTAAGTTGCCATTGCAAATCTTGTTGATGGTCATGAA
TTCCTTGGCTTTCCAAAACACCGTTATCGATTGGGTTAGAGATCACAGATTGCATCACAAGTA
CTCTGATACTGATGCTGATCCACATAATGCTTCTAGAGGTTTCTTCTACTCTCATGTTGGTTG
GTTGTTGGTTAGAAAACACCCAGATGTTAAGAAGAGAGGTAAAGAAATCGACATCTCCGACA
TCTACAACAACCCAGTTTTGAGATTCCAAAAGAAGTACGCCATTCCATTCATTGGTGCTGTTT
GTTTTGTTTTGCCAACCTTGATTCCAGTTTATGGTTGGGGTGAAACTTGGACTAATGCTTGGC
ATGTTGCTATGTTGAGATATATCATGAACTTGAACGTCACCTTCTTGGTTAATTCTGCTGCTC
ATATCTACGGTAAAAGACCATACGATAAGAAGATCTTGCCATCCCAAAACATTGCTGTTTCTA
TTGCTACTTTCGGTGAAGGTTTCCATAACTACCATCATGTTTTTCCATGGGATTACAGAGCTG
CTGAATTGGGTAACAATTCTTTGAACTTCCCTACCAAGTTCATCGATTTTTTCGCTTGGATTG
GTTGGGCCTACGATTTGAAAACTGTCTCCAAAGAAATGATCAAGCAAAGATCTAAGAGAACC

GGTGATGGTACTAATTTGTGGGGTTTGGAAGATGTTGATACCCCAGAAGATTTGAAGAACAC
TAAGGGTGAATGA

>SEQ ID NO: 11. Δ11-desaturase from *Trichoplusia ni*
ATGGCTGTGATGGCTCAAACAGTACAAGAAACGGCTACAGTGTTGGAAGAGGAAGCTCGCA
CAGTGACTCTTGTGGCTCCAAAGACAACGCCAAGGAAATATAAATATATATACACCAACTTTC
TTACATTTTCATATGCGCATTTAGCTGCATTATACGGACTTTATTTGTGCTTCACCTCTGCGAA
ATGGGAAACATTGCTATTCTCTTTCGTACTCTTCCACATGTCAAATATAGGCATCACCGCAGG
GGCTCACCGACTCTGGACTCACAAGACTTTCAAAGCCAAATTGCCTTTGGAAATTGTCCTCA
TGATATTCAACTCTTTAGCCTTTCAAAACACGGCTATTACATGGGCTAGAGAACATCGGCTAC
ATCACAAATACAGCGATACTGATGCTGATCCCCACAATGCGTCAAGAGGGTTCTTCTACTCG
CATGTTGGCTGGCTATTAGTAAAAAAACATCCCGATGTCCTGAAATATGGAAAAACTATAGAC
ATGTCGGATGTATACAATAATCCTGTGTTAAAATTTCAGAAAAAGTACGCAGTACCCTTAATT
GGAACAGTTTGTTTTGCTCTTCCAACTTTGATTCCAGTCTACTGTTGGGGCGAATCGTGGAA
CAACGCTTGGCACATAGCCTTATTTCGATACATATTCAATCTTAACGTGACTTTCCTAGTCAA
CAGTGCTGCGCATATCTGGGGGAATAAGCCTTATGATAAAAGCATCTTGCCCGCTCAAAACC
TGCTGGTTTCCTTCCTAGCAAGTGGAGAAGGCTTCCATAATTACCATCACGTCTTTCCATGG
GATTACCGCACAGCAGAATTAGGGAATAACTTCCTGAATTTGACGACGCTGTTCATTGATTTT
TGTGCCTGGTTTGGATGGGCTTATGACTTGAAGTCTGTATCAGAGGATATTATAAAACAGAG
AGCTAAACGAACAGGTGACGGTTCTTCAGGGGTCATTTGGGGATGGGACGACAAAGACATG
GACCGCGATATAAAATCTAAAGCTAACATTTTTTATGCTAAAAAGGAATGA

>SEQ ID NO: 12. Δ9-desaturase from *Drosophila melanogaster*
ATGGCTCCATACTCTAGAATCTACCACCAAGATAAGTCCTCTAGAGAAACTGGTGTTTTGTTC
GAAGATGATGCTCAAACCGTTGATTCTGATTTGACTACCGATAGATTCCAATTGAAGAGAGC
CGAAAAAAGAAGATTGCCATTGGTTTGGAGAAACATCATCTTGTTCGCTTTGGTTCATTTGGC
TGCCTTGTATGGTTTACATTCCATTTTCACTAGAGCTAAGTTGGCTACTACTTTGTTTGCTGCT
GGTTTGTACATTATCGGTATGTTGGGTGTTACTGCTGGTGCTCATAGATTGTGGGCTCATAG
AACTTACAAAGCTAAATGGCCTTTGAGATTGTTGTTGGTCATCTTCAACACCATTGCTTTCCA
AGATGCTGTTTATCATTGGGCCAGAGATCATAGAGTTCATCACAAATACTCTGAAACCGATG
CTGATCCACATAATGCTACTAGAGGTTTCTTCTTCTCTCATGTTGGTTGGTTGTTGTGCAAGA
AACACCCAGATATCAAAGAAAAGGGTAGAGGTTTGGATTTGTCCGATTTGAGAGCTGATCCA
ATCTTGATGTTTCAAAGAAAGCACTACTACATCTTGATGCCATTGGCTTGTTTTGTTTTGCCAA
CCGTTATTCCAATGGTCTACTGGAACGAAACTTTGGCTTCTTCTTGGTTTGTTGCTACTATGT
TCAGATGGTGCTTCCAATTGAATATGACCTGGTTGGTTAATTCCGCTGCTCATAAGTTTGGTA
ATAGACCATACGATAAGACCATGAACCCAACTCAAAATGCTTTCGTTTCTGCTTTCACTTTTG
GTGAAGGTTGGCATAATTACCATCATGCTTTTCCATGGGATTACAAGACTGCTGAATGGGGT
TGTTACTCTTTGAACATTACTACCGCCTTCATTGATTTGTTCGCTAAAATTGGTTGGGCCTAC
GATTTGAAAACTGTTGCTCCAGATGTTATCCAAAGAAGAGTTTTGAGAACTGGTGATGGTTCT
CATGAATTGTGGGGTTGGGGTGATAAGGATTTGACCGCTGAAGATGCTAGAAACGTTTTGTT
GGTTGACAAGTCCAGATAA

>SEQ ID NO: 13. Hygromycin resistance gene
ATGGGTAAAAAGCCTGAACTCACCGCGACGTCTGTCGAGAAGTTTCTGATCGAAAAGTTCGA
CAGCGTCTCCGACCTGATGCAGCTCTCGGAGGGCGAAGAATCTCGTGCTTTCAGCTTCGAT
GTAGGAGGGCGTGGATATGTCCTGCGGGTAAATAGCTGCGCCGATGGTTTCTACAAAGATC
GTTATGTTTATCGGCACTTTGCATCGGCCGCGCTCCCGATTCCGGAAGTGCTTGACATTGGG
GAATTCAGCGAGAGCCTGACCTATTGCATCTCCCGCCGTGCACAGGGTGTCACGTTGCAAG
ACCTGCCTGAAACCGAACTGCCCGCTGTTCTGCAGCCGGTCGCGGAGGCAATGGATGCCAT
TGCTGCGGCCGATCTTAGCCAGACGAGCGGGTTCGGCCCATTCGGACCGCAAGGAATCGG
TCAATACACTACATGGCGTGATTTCATATGCGCGATTGCTGATCCCCATGTGTATCACTGGC
AAACTGTGATGGACGACACCGTCAGTGCGTCCGTCGCGCAGGCTCTCGATGAGCTGATGCT
TTGGGCCGAGGACTGCCCCGAAGTCCGGCACCTCGTGCACGCGGATTTCGGCTCCAACAA
TGTCCTGACGGACAATGGCCGCATAACAGCGGTCATTGACTGGAGCGAGGCGATGTTCGG
GGATTCCCAATACGAGGTCGCCAACATCTTCTTCTGGAGGCCGTGGTTGGCTTGTATGGAG
CAGCAGACGCGCTACTTCGAGCGGAGGCATCCGGAGCTTGCAGGATCGCCGCGGCTCCGG

GCGTATATGCTCCGCATTGGTCTTGACCAACTCTATCAGAGCTTGGTTGACGGCAATTTCGA
TGATGCAGCTTGGGCGCAGGGTCGATGCGACGCAATCGTCCGATCCGGAGCCGGGACTGT
CGGGCGTACACAAATCGCCCGCAGAAGCGCGGCCGTCTGGACCGATGGCTGTGTAGAAGT
ACTCGCCGATAGTGGAAACCGACGCCCCAGCACTCGTCCGAGGGCAAAGGAATAA

>SEQ ID NO: 14. Atrd11 expression cassette and the upstream genomic region of integration site B

| Element | Position (bp) |
|---|---|
| IntB upstream | 9-508 |
| PEX20 terminator | 881-569 |
| Atrd11 | 1862-882 |
| GPD promoter | 2800-1869 |

CGTGCGATCCCACAGTTCTCACTCAGATCATGGAGACTCTAACCTTGAGACATCAATTATCA
GCTCTCGAGGATAATGTTAGTGCAGTTCCAGGACTCATTGTGCAACTGTCACCACGGCATTT
TGGGTCTGTTCTTTTGAAGTACAGAAATATCCTCATTGTTGGTATACTTTGGGACTTTTTCTTG
TTACAGAAGAATAAAAAAACCTCGACTGATGTACTAATTACATGGTTAACATCCCCAAGGTCA
AAGTACAGATATTGTACCGACTTCTGAAATTTGTGGGATCCACACACGGACCTCTGCGATGA
TACAATATCATGGTTCCATGGTCTCTTGAATCACACCACTCAATAATAAACACCAGCTCTTTC
AATCCAACTTAGCTTTCTTCCACTTGAACATAGGCCATCCCCCCTTCTTTCTATTTTACATAAA
TAGCAAGATCCTTACCCCTACATGTCTCATAACACAATCTCAACTTGACTTCCCATAAGAAGT
TCACTCAGTCATGAAAGTCTTAGTACTGGACGTGCAACGCTTCAGATGTGACCATATACTTA
GGCACGCAACTAACATGAATGAATACGATATACATCAAAGACTATGATACGCAGTATTGCAC
ACTGTACGAGTAAGAGCACTAGCCACTGCACTCAAGTGAAACCGTTGCCCGGGTACGAGTA
TGAGTATGTACAGTATGTTTAGTATTGTACTTGGACAGTGCTTGTATCGTACATTCTCAAGTG
TCAAACATAAATATCCGTTGCTATATCCTCGCACCACCACGTAGCTCGCTATATCCCTGTGTT
GAATCCATCCATCTTGGATTGCCAATTGTGCACACAGAACCGGGCACTCACTTCCCCATCCA
CACTTTCATCTCTTTCTACCCCACATATCAGTACCATCACCAGTTTTAGCACATCTCTTAGCAA
TAGCTTCTTGTGGAACAGATTTCAAATCGTAGGCCCAACCAATCCAAGCAAAGAAATCAATG
AAGGCGGTGGTCATATTCAATCTATTGTTACCCAATTCAGCAGCTCTGTAATCCCATGGATAA
GTATGATGGTAGTTATGGAAACCTTCACCGAAAGTAGCCAAAGAAACAGAGATGTTTTGAAC
AGAGGCGATAGACTTATCGTATGGCTTGTTACCAAAAATATGAGCAGCGGAGTTAACCAAAA
AGGTCATGTTCAAGTTCATGACGTATCTTAACAAGTTAATGTTCCAGGCAGTAGAAATACCTT
CACCCCAAAAGTAAACTGGAACGAAAGTTGGCATCAAAAAGCACAAAGTACCAATGACCAAA
ATGGCGTACTTCTTTTGGAACTTCAACAAAGGGTTGTTCTTCAAGTCATCCAAAGACAAGTAC
TTACCTCTAGCCTTAACTTCTGGGTGTTTCTTAACCAACAACCAACCAACATGAGAGTAGAAG
AAACCTCTAGTAGCATTATGTGGATCAGCATCAGTATCAGAGTACTTGTGATGCAATCTATGA
TCTCTAGCCCAAGTAAAAGCAGTATCTTGGAAGGCAATGGAGTTCATGATCAACAACAAGAT
TTCCAATGGCAACTTGGCTTTGTAAGTTCTATGAGCCCATAATCTATGAGCACCACCAGTAAT
ACCAATTTCGGCGATAACGTACAAGAAGAAAGCGAACAAAATAGTAGCCCACTTAGCACAAG
TGAAACACAAGTACAAACCATAAACAGCAGACAAATGCCAGTAACCGAAGGTCAACAAGTTT
CTGTAAACGATTTTGTACTTTCTTGGACCAGCTTGTGGAGCAATCAACTTGGTGAATTGTGGT
TCAGAATGTTCAGACAAAACATCAGAGGAACCCTTGTTTGGAACCATTGTGGCGTTGATGTG
TGTTTAATTCAAGAATGAATATAGAGAAGAGAAGAAGAAAAAAGATTCAATTGAGCCGGCGAT
GCAGACCCTTATATAAATGTTGCCTTGGACAGACGGAGCAAGCCCGCCCAAACCTACGTTC
GGTATAATATGTTAAGCTTTTTAACACAAAGGTTTGGCTTGGGGTAACCTGATGTGGTGCAAA
AGACCGGGCGTTGGCGAGCCATTGCGCGGGCGAATGGGGCCGTGACTCGTCTCAAATTCG
AGGGCGTGCCTCAATTCGTGCCCCCGTGGCTTTTTCCCGCCGTTTCCGCCCCGTTTGCACC
ACTGCAGCCGCTTCTTTGGTTCGGACACCTTGCTGCGAGCTAGGTGCCTTGTGCTACTTAAA
AAGTGGCCTCCCAACACCAACATGACATGAGTGCGTGGGCCAAGACACGTTGGCGGGGTC
GCAGTCGGCTCAATGGCCCGGAAAAAACGCTGCTGGAGCTGGTTCGGACGCAGTCCGCCG
CGGCGTATCGATATCCGCAAGGTTCCATGGCGCCATTGCCCTCCGTCGGCGTCTATCCCGC
AACCTCTAAATAGAGCGGGAATATAACCCAAGCTTCTTTTTTTTTCCTTTAACACGCACACCC
CCAACTATCATGTTGCTGCTGCTGTTTGACTCTACTCTGTGGAGGGGTGCTCCCACCCAACC
CAACCTACAGGTGGATCCGGCGCTGTGATTGGCTGATAAGTCTCCTATCCGGACTAATTCTG

ACCAATGGGACATGCGCGCAGGACCCAAATGCCGCAATTACGTAACCCCAACGAAATGCCT
ACCCCTCTTTGGAGCCCAGCGGCCCCAAATCCCCCCAAGCAGCCCGGTTCTACCGGCTTCC
ATCTCCAAGCACCCCTTTCTCCACACCCCACAAAAAGACCCGTGCAGGACATCCTACTGCGT
CACCTGCACT

>SEQ ID NO: 15. HsuFAR expression cassette

| Element | Position (bp) |
|---|---|
| TEFINTRON promoter | 10-540 |
| HsuFAR | 541-1899 |
| LIP2 terminator | 1900-2836 |

ACCTGCACTAGAGACCGGGTTGGCGGCGCATTTGTGTCCCAAAAAACAGCCCCAATTGCCC
CAATTGACCCCAAATTGACCCAGTAGCGGGCCCAACCCCGGCGAGAGCCCCCTTCTCCCCA
CATATCAAACCTCCCCCGGTTCCCACACTTGCCGTTAAGGGCGTAGGGTACTGCAGTCTGG
AATCTACGCTTGTTCAGACTTTGTACTAGTTTCTTTGTCTGGCCATCCGGGTAACCCATGCCG
GACGCAAAATAGACTACTGAAAATTTTTTTGCTTTGTGGTTGGGACTTTAGCCAAGGGTATAA
AAGACCACCGTCCCCGAATTACCTTTCCTCTTCTTTTCTCTCTCTCCTTGTCAACTCACACCC
GAAATCGTTAAGCATTTCCTTCTGAGTATAAGAATCATTCAAAATGGTGAGTTTCAGAGGCAG
CAGCAATTGCCACGGGCTTTGAGCACACGGCCGGGTGTGGTCCCATTCCCATCGACACAAG
ACGCCACGTCATCCGACCAGCACTTTTTGCAGTACTAACCGCAGGTTGTCTTGACCTCCAAA
GAAACTAAGCCATCTGTTGCTGAATTTTACGCTGGTAAGTCTGTTTTCATTACTGGTGGTACT
GGTTTCTTGGGTAAGGTTTTCATTGAAAAGTTGTTGTACTCCTGCCCAGATATCGGTAATATC
TACATGTTGATCAGAGAAAAGAAGGGTTTGTCCGTTTCCGAAAGAATCAAGCACTTTTTGGAT
GATCCTTTGTTCACCAGATTGAAAGAAAAAAGACCAGCCGACTTGGAAAAGATCGTTTTGATT
CCAGGTGATATTACTGCTCCAGATTTGGGTATTACCTCCGAAAACGAAAAGATGTTGATCGA
AAAGGTCAGTGTCATTATTCATTCTGCTGCTACCGTTAAGTTCAACGAACCATTGCCAACTGC
TTGGAAGATTAACGTTGAAGGTACTAGAATGATGTTGGCCTTGTCTAGAAGAATGAAGAGAA
TCGAAGTTTTCATCCATATCTCTACCGCTTACACTAACACCAACAGAGAAGTTGTTGACGAAA
TCTTGTATCCAGCTCCAGCTGATATTGATCAAGTTCACCAATATGTTAAGGACGGTATCTCTG
AAGAAGAAACTGAAAAAATCTTGAACGGTAGACCAAACACTTACACTTTCACTAAGGCTTTGA
CCGAACATTTGGTTGCTGAAAATCAAGCTTACGTTCCAACCATTATCGTTAGACCATCAGTTG
TTGCTGCCATTAAGGATGAACCTATTAAGGGTTGGTTGGGTAATTGGTATGGTGCTACAGGT
TTGACTGTTTTTACTGCTAAGGGTTTGAACAGAGTTATCTACGGTCACTCTTCTAACATCGTT
GATTTGATCCCAGTTGATTACGTTGCCAACTTGGTTATTGCTGCTGGTGCTAAATCTTCTAAG
TCTACTGAATTGAAGGTCTACAACTGCTGTTCTTCTGCTTGTAACCCAATTACTATCGGTAAG
TTGATGTCCATGTTTGCTGAAGATGCTATCAAGCAAAAGTCTTACGCTATGCCATTGCCAGGT
TGGTACATTTTTACTAAGTACAAGTGGTTGGTCTTGTTGTTGACCATTTTGTTCCAAGTTATTC
CAGCCTACATTACCGACTTGTACAGACATTTGATTGGTAAGAACCCAAGATATATCAAGTTGC
AATCCTTGGTCAATCAAACCAGATCCTCCATTGATTTCTTCACCAACCATTCTTGGGTTATGA
AGGCTGATAGAGTCAGAGAATTATTCGCTTCTTTGTCTCCAGCAGATAAGTACTTGTTTCCAT
GTGATCCAGTCAACATCAATTGGAGACAATATATCCAAGATTACTGCTGGGGTGTTAGACATT
TCTTGGAAAAAAAGACTTAACTTCTGTTCGGAATCAACCTCAAGGTTAACGGCCACGATCCC
CTCGTTGTTACTCTTGGTCAGCCCATTGTCGGTAACGCTGGCTTTGCTAACTGGGTCGATAA
ACTCTTCTTTGGCCAGGAGAACCCCGATGTCTCCAAGGTGTCCAAAGACCGAAAGCTCTACC
GAATCACCCACCGAGGAGATATCGTCCCTCAAGTGCCCTTCTGGGACGGTTACCAGCACTG
CTCTGGTGAGGTCTTTATTGACTGGCCCCTGATCCACCCTCCTCTCTCCAACGTTGTCATGT
GCCAGGGCCAGAGCAATAAACAGTGCTCTGCCGGTAACACTCTGCTCCAGCAGGTCAATGT
GATTGGAAACCATCTGCAGTACTTCGTCACCGAGGGTGTCTGTGGTATCTAAGCTATTTATC
ACTCTTTACAACTTCTACCTCAACTATCTACTTTAATAAATGAATATCGTTTATTCTCTATGATT
ACTGTATATGCGTTCCTCTAAGACAAATCGAAACCAGCATGCGATCGAATGGCATACAAAAG
TTTCTTCCGAAGTTGATCAATGTCCTGATAGTCAGGCAGCTTGAGAAGATTGACACAGGTGG
AGGCCGTAGGGAACCGATCAACCTGTCTACCAGCGTTACGAATGGCAAATGACGGGTTCAA
AGCCTTGAATCCTTGCAATGGTGCCTTGGATACTGATGTCACAAACTTAAGAAGCAGCCGCT
TGTCCTCTTCCTCGAAACTCTCAAACACAGTCCAGAAGTCCTTTATAGTTTGATCTGTATCCA
GATAGCCTCCGTAATTGGTGTGTGTCTTCAAATCCCAGACGTCCACATTGGCATGTCCTCCA

CTGATAAGCATTTGAAGTTCATCTGCGTTGAACATTGAGACCCACGAAGGGTCAATGAGCTG
GTATAGACCGCCCAAGAATGCATCTGATCTGTCAT

>SEQ ID NO: 16. Ura3 marker cassette fused to 500 bp downstream region for integration into IntB site

| Element | Position (bp) |
|---|---|
| loxP site | 1-34 |
| EXP promoter | 35-1036 |
| Ura3 | 1037-1891 |
| *Cyc1* terminator | 1892-2124 |
| loxP site | 2125-2158 |
| IntB downstream | 2219-2704 |

ATAACTTCGTATAATGTATGCTATACGAAGTTATAAGGAGTTTGGCGCCCGTTTTTTCGAGCC
CCACACGTTTCGGTGAGTATGAGCGGCGGCAGATTCGAGCGTTTCCGGTTTCCGCGGCTGG
ACGAGAGCCCATGATGGGGGCTCCCACCACCAGCAATCAGGGCCCTGATTACACACCCACC
TGTAATGTCATGCTGTTCATCGTGGTTAATGCTGCTGTGTGCTGTGTGTGTGTGTTGTTTGG
CGCTCATTGTTGCGTTATGCAGCGTACACCACAATATTGGAAGCTTATTAGCCTTTCTATTTT
TTCGTTTGCAAGGCTTAACAACATTGCTGTGGAGAGGGATGGGGATATGGAGGCCGCTGGA
GGGAGTCGGAGAGGCGTTTTGGAGCGGCTTGGCCTGGCGCCCAGCTCGCGAAACGCACCT
AGGACCCTTTGGCACGCCGAAATGTGCCACTTTTCAGTCTAGTAACGCCTTACCTACGTCAT
TCCATGCATGCATGTTTGCGCCTTTTTTCCCTTGCCCTTGATCGCCACACAGTACAGTGCAC
TGTACAGTGGAGGTTTTGGGGGGGTCTTAGATGGGAGCTAAAAGCGGCCTAGCGGTACACT
AGTGGGATTGTATGGAGTGGCATGGAGCCTAGGTGGAGCCTGACAGGACGCACGACCGGC
TAGCCCGTGACAGACGATGGGTGGCTCCTGTTGTCCACCGCGTACAAATGTTTGGGCCAAA
GTCTTGTCAGCCTTGCTTGCGAACCTAATTCCCAATTTTGTCACTTCGCACCCCCATTGATCG
AGCCCTAACCCCTGCCCATCAGGCAATCCAATTAAGCTCGCATTGTCTGCCTTGTTTAGTTT
GGCTCCTGCCCGTTTCGGCGTCCACTTGCACAAACACAAACAAGCATTATATATAAGGCTCG
TCTCTCCCTCCCAACCACACTCACTTTTTTGCCCGTCTTCCCTTGCTAACACAAAAGTCAAGA
ACACAAACAACCACCCCAACCCCCTTACACACAAGACATATCTACAGCAATGCCCTCCTACG
AGGCCCGAGCCAACGTCCACAAGTCCGCCTTCGCCGCCCGAGTCCTGAAGCTGGTCGCCG
CCAAGAAGACCAACCTGTGCGCCTCCCTGGACGTCACCACCACCAAGGAGCTGATCGAGCT
GGCCGACAAGGTCGGCCCCTACGTCTGCATGATCAAGACCCACATCGACATCATCGACGAC
TTCACCTACGCCGGCACCGTCCTGCCCCTGAAGGAGCTGGCCCTGAAGCACGGCTTCTTCC
TGTTCGAGGACCGAAAGTTCGCCGACATCGGCAACACCGTCAAGCACCAGTACCGATGCCA
CCGAATCGCCGAGTGGTCCGACATCACCAACGCCCACGGCGTCCCCGGCACCGGCATCAT
CGCCGGCCTGCGAGCCGGCGCCGAGGAGACCGTCTCCGAGCAGAAGAAGGAGGACGTCT
CCGACTACGAGAACTCCCAGTACAAGGAGTTCCTGGTCCCCTCCCCCAACGAGAAGCTGGC
CCGAGGCCTGCTGATGCTGGCCGAGCTGTCCTGCAAGGGCTCCCTGGCCACCGGCGAGTA
CTCCAAGCAGACCATCGAGCTGGCCCGATCCGACCCCGAGTTCGTCGTCGGCTTCATCGCC
CAGAACCGACCCAAGGGCGACTCCGAGGACTGGCTGATCCTGACCCCCGGCGTCGGCCTG
GACGACAAGGGCGACGCCCTGGGCCAGCAGTACCGAACCGTCGAGGACGTCATGTCCACC
GGCACCGACATCATCATCGTCGGCCGAGGCCTGTACGGCCAGAACCGAGACCCCATCGAG
GAGGCCAAGCGATACCAGAAGGCCGGCTGGGAGGCCTACCAGAAGATCAACTGCTAGTCA
TGTAATTAGTTATGTCACGCTTACATTCACGCCCTCCCCCCACATCCGCTCTAACCGAAAAG
GAAGGAGTTAGACAACCTGAAGTCTAGGTCCCTATTTATTTTTTTATAGTTATGTTAGTATTAA
GAACGTTATTTATATTTCAAATTTTTCTTTTTTTTCTGTACAGACGCGTGTACGCATGTAACAT
TATACTGAAAACCTTGCTTGAGAAGGTTTTGGGACGCTCGATAACTTCGTATAATGTATGCTA
TACGAAGTTATCAGCATCGTAATAGCCTCCAAGAGATTGATCATCACTCTGAATGTACAAGCA
ACCCAAGTACCTGCTCCTGCACCTAAGTTCGTCAAAATCGGTTTTACTCAGAGAATCAACAA
CCCTACCAACTGTACATACTGCTAACCCTGATTCTTTGAATAACCCCAATAAGGCTCCTGCTG
ACCCTTCTGCCGTTCTAGGAAACCAGCTTGTTGAGTACGCTGTGAACATCACTCTTGGTATA
CCCGGTCAGCCCTTCTCAGTTCAGATTGACACAGGCTCGTCTGACTTGTGGGTGAAGAGTG
ACGGCTCCTCCGGTGCATTCAACAAGAAGGCTTCTTCTACTTTTCAGGAGGACGTTCCCAAC
GGCTTTGCAATTGCCCACGGAGACAAAACCTCTGCCATTGGAGATTGGGTCAAAGATACCAT

CAATATCGGTGGTGTTAGTATTGACCAGTATAGTATTGACCAGTATGAGTTCGCCATGGCTA
CTCAGACAAATACTGACCCGGTTTTTGGTATCGGCTACCCGAGCAACGAGGCGTCTTATG

>SEQ ID NO: 17. Atrd11 expression cassette

| Element | Position (bp) |
|---|---|
| PEX20 terminator | 321-9 |
| Atrd11 | 1302-322 |
| GPD promoter | 2240-1309 |

CGTGCGATACGCAACTAACATGAATGAATACGATATACATCAAAGACTATGATACGCAGTATT
GCACACTGTACGAGTAAGAGCACTAGCCACTGCACTCAAGTGAAACCGTTGCCCGGGTACG
AGTATGAGTATGTACAGTATGTTTAGTATTGTACTTGGACAGTGCTTGTATCGTACATTCTCA
AGTGTCAAACATAAATATCCGTTGCTATATCCTCGCACCACCACGTAGCTCGCTATATCCCTG
TGTTGAATCCATCCATCTTGGATTGCCAATTGTGCACACAGAACCGGGCACTCACTTCCCCA
TCCACACTTTCATCTCTTTCTACCCCACATATCAGTACCATCACCAGTTTTAGCACATCTCTTA
GCAATAGCTTCTTGTGGAACAGATTTCAAATCGTAGGCCCAACCAATCCAAGCAAAGAAATC
AATGAAGGCGGTGGTCATATTCAATCTATTGTTACCCAATTCAGCAGCTCTGTAATCCCATGG
ATAAGTATGATGGTAGTTATGGAAACCTTCACCGAAAGTAGCCAAAGAAACAGAGATGTTTT
GAACAGAGGCGATAGACTTATCGTATGGCTTGTTACCAAAAATATGAGCAGCGGAGTTAACC
AAAAAGGTCATGTTCAAGTTCATGACGTATCTTAACAAGTTAATGTTCCAGGCAGTAGAAATA
CCTTCACCCCAAAAGTAAACTGGAACGAAAGTTGGCATCAAAAAGCACAAAGTACCAATGAC
CAAAATGGCGTACTTCTTTTGGAACTTCAACAAAGGGTTGTTCTTCAAGTCATCCAAAGACAA
GTACTTACCTCTAGCCTTAACTTCTGGGTGTTTCTTAACCAACAACCAACCAACATGAGAGTA
GAAGAAACCTCTAGTAGCATTATGTGGATCAGCATCAGTATCAGAGTACTTGTGATGCAATCT
ATGATCTCTAGCCCAAGTAAAAGCAGTATCTTGGAAGGCAATGGAGTTCATGATCAACAACA
AGATTTCCAATGGCAACTTGGCTTTGTAAGTTCTATGAGCCCATAATCTATGAGCACCACCAG
TAATACCAATTTCGGCGATAACGTACAAGAAGAAAGCGAACAAAATAGTAGCCCACTTAGCA
CAAGTGAAACACAAGTACAAACCATAAACAGCAGACAAATGCCAGTAACCGAAGGTCAACAA
GTTTCTGTAAACGATTTTGTACTTTCTTGGACCAGCTTGTGGAGCAATCAACTTGGTGAATTG
TGGTTCAGAATGTTCAGACAAAACATCAGAGGAACCCTTGTTTGGAACCATTGTGGCGTTGA
TGTGTGTTTAATTCAAGAATGAATATAGAGAAGAGAAGAAGAAAAAAGATTCAATTGAGCCGG
CGATGCAGACCCTTATATAAATGTTGCCTTGGACAGACGGAGCAAGCCCGCCCAAACCTAC
GTTCGGTATAATATGTTAAGCTTTTTAACACAAAGGTTTGGCTTGGGGTAACCTGATGTGGTG
CAAAAGACCGGGCGTTGGCGAGCCATTGCGCGGGCGAATGGGGCCGTGACTCGTCTCAAA
TTCGAGGGCGTGCCTCAATTCGTGCCCCCGTGGCTTTTTCCCGCCGTTTCCGCCCCGTTTG
CACCACTGCAGCCGCTTCTTTGGTTCGGACACCTTGCTGCGAGCTAGGTGCCTTGTGCTAC
TTAAAAAGTGGCCTCCCAACACCAACATGACATGAGTGCGTGGGCCAAGACACGTTGGCGG
GGTCGCAGTCGGCTCAATGGCCCGGAAAAAACGCTGCTGGAGCTGGTTCGGACGCAGTCC
GCCGCGGCGTATCGATATCCGCAAGGTTCCATGGCGCCATTGCCCTCCGTCGGCGTCTATC
CCGCAACCTCTAAATAGAGCGGGAATATAACCCAAGCTTCTTTTTTTTTCCTTTAACACGCAC
ACCCCCAACTATCATGTTGCTGCTGCTGTTTGACTCTACTCTGTGGAGGGGTGCTCCCACCC
AACCCAACCTACAGGTGGATCCGGCGCTGTGATTGGCTGATAAGTCTCCTATCCGGACTAAT
TCTGACCAATGGGACATGCGCGCAGGACCCAAATGCCGCAATTACGTAACCCCAACGAAAT
GCCTACCCCTCTTTGGAGCCCAGCGGCCCCAAATCCCCCCAAGCAGCCCGGTTCTACCGG
CTTCCATCTCCAAGCACCCCTTTCTCCACACCCCACAAAAAGACCCGTGCAGGACATCCTAC
TGCGTCACCTGCACT

>SEQ ID NO: 18. *LEU2* from *Kluyveromyces lactis*
ATGTCTAAGAATATCGTTGTCCTACCGGGTGATCACGTCGGTAAAGAAGTTACTGACGAAGC
TATTAAGGTCTTGAATGCCATTGCTGAAGTCCGTCCAGAAATTAAGTTCAATTTCCAACATCA
CTTGATCGGGGGTGCTGCCATCGATGCCACTGGCACTCCTTTACCAGATGAAGCTCTAGAA
GCCTCTAAGAAAGCCGATGCTGTCTTACTAGGTGCTGTTGGTGGTCCAAAATGGGGTACGG
GCGCAGTTAGACCAGAACAAGGTCTATTGAAGATCAGAAAGGAATTGGGTCTATACGCCAAC
TTAAGACCATGTAACTTTGCTTCTGATTCTTTACTAGATCTTTCTCCTTTGAAGCCTGAATATG
CAAAGGGTACCGATTTCGTCGTCGTTAGAGAATTGGTTGGTGGTATCTACTTTGGTGAAAGA
AAAGAAGATGAAGGTGACGGAGTTGCTTGGGACTCTGAGAAATACAGTGTTCCTGAAGTTCA

AAGAATTACAAGAATGGCTGCTTTCTTGGCATTGCAACAAAACCCACCATTACCAATCTGGTC
ACTTGACAAGGCTAACGTGCTTGCCTCTTCCAGATTGTGGAGAAAGACTGTTGAAGAAACCA
TCAAGACTGAGTTCCCACAATTAACTGTTCAGCACCAATTGATCGACTCTGCTGCTATGATTT
TGGTTAAATCACCAACTAAGCTAAACGGTGTTGTTATTACCAACAACATGTTTGGTGATATTA
TCTCCGATGAAGCCTCTGTTATTCCAGGTTCTTTGGGTTTATTACCTTCTGCATCTCTAGCTT
CCCTACCTGACACTAACAAGGCATTCGGTTTGTACGAACCATGTCATGGTTCTGCCCCAGAT
TTACCAGCAAACAAGGTTAACCCAATTGCTACCATCTTATCTGCAGCTATGATGTTGAAGTTA
TCCTTGGATTTGGTTGAAGAAGGTAGGGCTCTTGAAGAAGCTGTTAGAAATGTCTTGGATGC
AGGTGTCAGAACCGGTGACCTTGGTGGTTCTAACTCTACCACTGAGGTTGGCGATGCTATC
GCCAAGGCTGTCAAGGAAATCTTGGCTTAA

>SEQ ID NO: 19. Fatty acyl-reductase from *Helicoverpa armigera* codon-optimized for *Y. lipolytica*
ATGGTGGTCCTGACCTCTAAGGAGACTAAGCCCTCCGTGGCCGAGTTCTACGCTGGCAAGT
CTGTCTTCATCACCGGCGGAACCGGTTTCCTGGGCAAGGTCTTCATTGAGAAGCTGCTGTA
CTCCTGTCCCGACATCGGCAACATCTACATGCTGATCCGAGAGAAGAAGGGACTGTCTGTG
TCCGAGCGAATTAAGCACTTCCTGGACGACCCCCTGTTCACCCGACTGAAGGAGAAGCGAC
CCGCCGACCTGGAGAAGATCGTGCTGATTCCCGGAGACATCACCGCTCCCGACCTGGGTAT
TACCTCTGAGAACGAGAAGATGCTGATCGAGAAGGTGTCTGTCATCATTCACTCCGCCGCTA
CCGTCAAGTTCAACGAGCCCCTGCCCACCGCCTGGAAGATCAACGTGGAGGGAACCCGAAT
GATGCTGGCTCTGTCTCGACGAATGAAGCGAATTGAGGTCTTCATCCACATTTCCACCGCCT
ACACCAACACCAACCGAGAGGTGGTGGACGAGATCCTGTACCCTGCTCCTGCTGACATTGA
CCAGGTGCACCGATACGTCAAGGACGGTATCTCTGAGGAAGAGACTGAGAAGATTCTGAAC
GGCCGACCCAACACCTACACCTTCACCAAGGCCCTGACCGAGCACCTGGTGGCTGAGAAC
CAGGCTTACGTGCCCACCATCATTGTCCGACCCTCCGTGGTCGCCGCTATCAAGGACGAGC
CCATTAAGGGATGGCTGGGTAACTGGTACGGAGCTACCGGACTGACCGTGTTCACCGCTAA
GGGTCTGAACCGAGTCATCTACGGCCACTCTTCCAACATCGTGGACCTGATTCCCGTGGAC
TACGTCGCCAACCTGGTCATTGCCGCTGGCGCTAAGTCTTCCAAGTCCACCGAGCTGAAGG
TGTACAACTGTTGCTCTTCCGCCTGCAACCCCATCACCATTGGAAAGCTGATGTCTATGTTC
GCCGAGGACGCTATCAAGCAGAAGTCCTACGCTATGCCCCTGCCCGGTTGGTACATCTTCA
CCAAGTACAAGTGGCTGGTCCTGCTGCTGACCATTCTGTTCCAGGTCATCCCCGCCTACATT
ACCGACCTGTACCGACACCTGATCGGCAAGAACCCCCGATACATTAAGCTGCAGTCTCTGG
TCAACCAGACCCGATCTTCCATTGACTTCTTCACCTCTCACTCCTGGGTCATGAAGGCTGAC
CGAGTCCGAGAGCTGTTCGCCTCTCTGTCCCCCGCTGACAAGTACCTGTTCCCCTGTGACC
CCACCGACATCAACTGGACCCACTACATTCAGGACTACTGCTGGGGAGTGCGACACTTCCT
GGAGAAGAAGTCCTACGAGTAG

>SEQ ID NO: 20. Δ9-desaturase from *Drosophila melanogaster* codon.optimized for *Y. lipolytica*
ATGGCTCCCTACTCTCGAATCTACCACCAGGACAAGTCGTCCCGAGAGACTGGCGTGCTGT
TCGAGGACGACGCCCAGACCGTGGACTCTGACCTGACCACCGACCGATTCCAGCTGAAGC
GAGCCGAGAAGCGACGACTGCCCCTGGTGTGGCGAAACATCATCCTGTTCGCCCTGGTGC
ACCTGGCCGCTCTGTACGGCCTGCACTCTATCTTCACCCGAGCCAAGCTGGCCACCACTCT
GTTCGCTGCCGGCCTGTACATCATCGGCATGCTGGGCGTGACCGCTGGCGCCCACCGACT
GTGGGCTCACCGAACCTACAAGGCCAAGTGGCCCCTGCGACTGCTGCTGGTGATCTTCAAC
ACCATTGCCTTCCAGGACGCCGTGTACCACTGGGCCCGAGATCACCGAGTGCACCACAAGT
ACTCTGAGACTGACGCTGACCCTCACAACGCTACCCGAGGCTTCTTCTTCTCTCACGTCGGC
TGGCTGCTGTGCAAGAAGCACCCCGACATCAAGGAAAAGGGCCGAGGCCTGGACCTGTCT
GACCTGCGAGCTGACCCCATCCTGATGTTCCAGCGAAAGCACTACTACATTCTGATGCCCCT
GGCCTGCTTCGTGCTGCCCACCGTGATTCCCATGGTGTACTGGAACGAGACTCTGGCCTCT
TCCTGGTTCGTGGCCACCATGTTCCGATGGTGCTTCCAGCTCAACATGACCTGGCTGGTGA
ACTCTGCCGCTCACAAGTTCGGCAACCGACCTTACGACAAGACTATGAACCCCACTCAGAAC
GCCTTCGTGTCTGCCTTCACCTTCGGCGAAGGCTGGCACAACTACCACCACGCATTCCCTT
GGGACTACAAGACCGCCGAGTGGGGCTGCTACTCTCTGAACATCACCACCGCCTTCATCGA
CCTGTTCGCTAAGATCGGCTGGGCCTACGACCTCAAGACCGTGGCTCCCGACGTGATCCAG
CGACGAGTGCTGCGAACCGGCGACGGCTCTCACGAGCTGTGGGGCTGGGGCGACAAGGA
CCTGACCGCTGAGGACGCCCGAAACGTCCTGCTGGTGGACAAGTCTCGATAA

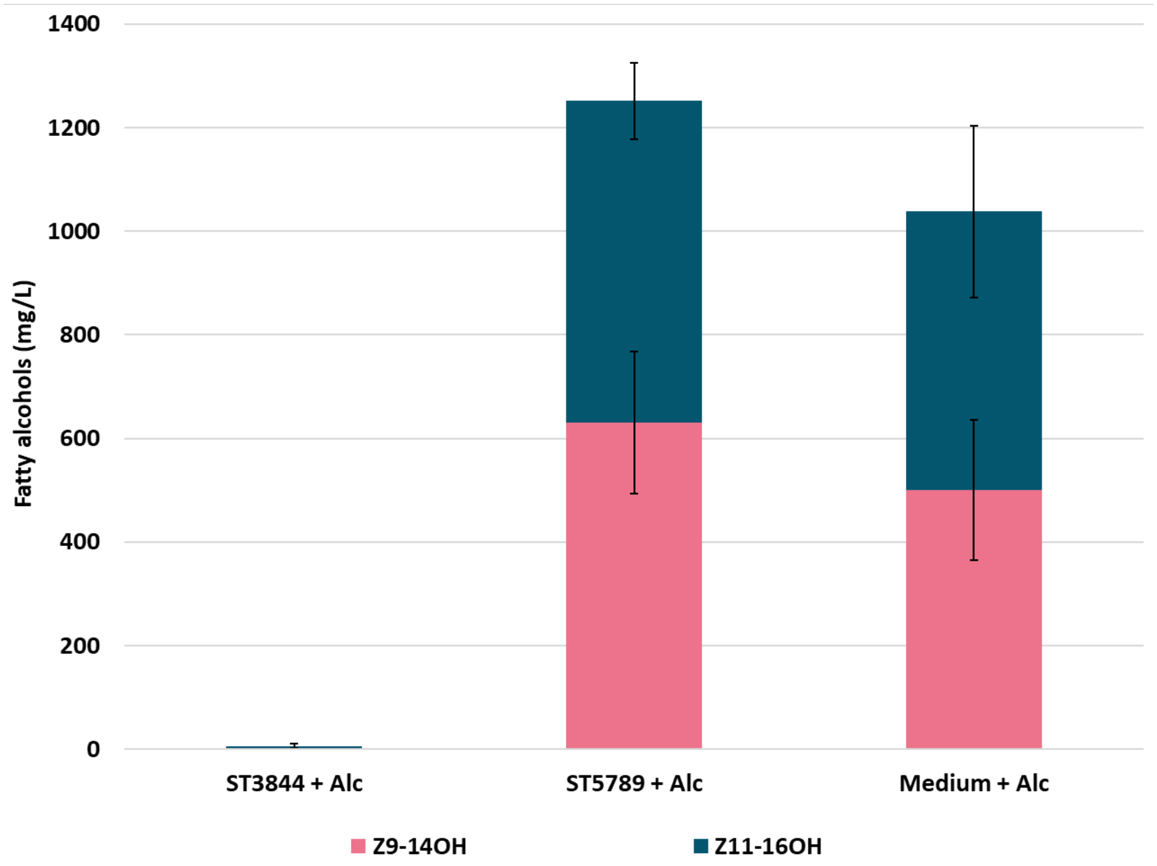


**Fig. S1.** Analysis of external fatty alcohol degradation in a *Y. lipolytica* strain expressing the heterologous genes HsFAR and Atrd11, and being devoid of the intrinsic genes *HFD1*/*HFD4*/*FAO1*/*PEX10* (ST5789), as well as in a strain only expressing the heterologous genes HsFAR and Atrd11 (ST3844). As a control, external fatty alcohols were added to culture medium (medium + alc) and the alcohol concentration was determined after the same incubation time.

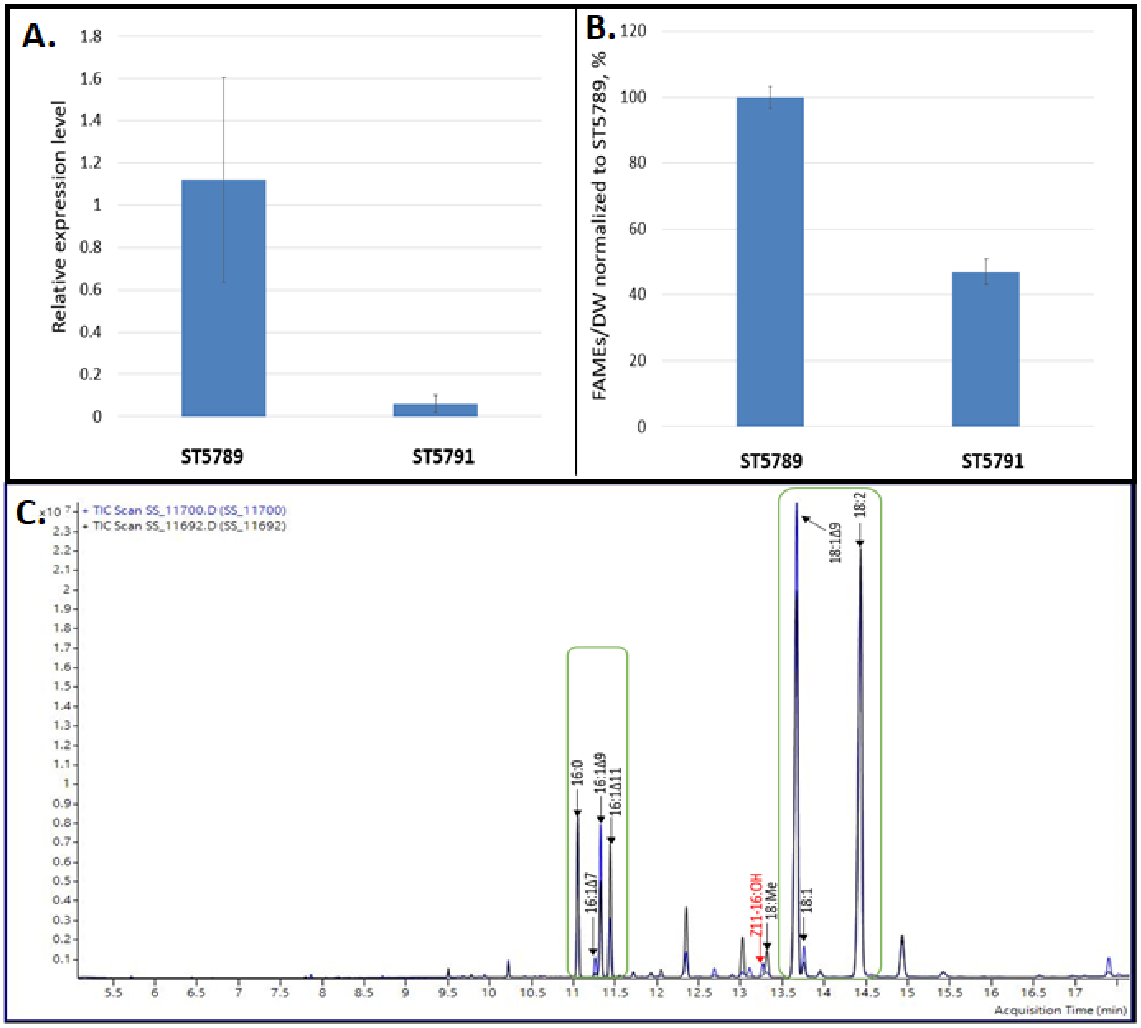


**Fig. S2.** Expression of glycerol-3-phosphate acyltransferase gene and fatty acid content in the strain with truncated GPAT promoter. **A.** Expression of *GPAT* gene in the control strain (ST5789) and in a strain with truncated *GPAT* promoter (ST5791) measured by qRT-PCR. **B.** Total fatty acid content measured as fatty acid methyl esters. **C.** Overlaid chromatograms of FAMEs extracts from ST5789 (black) and ST5791 (blue) strains. Green circles indicate the methyl esters that were quantified and included in total FAMEs content in panel B. Methyl octadecanoate (18Me) was not included in the FAMEs calculation because it co-eluted with Z11-16OH.

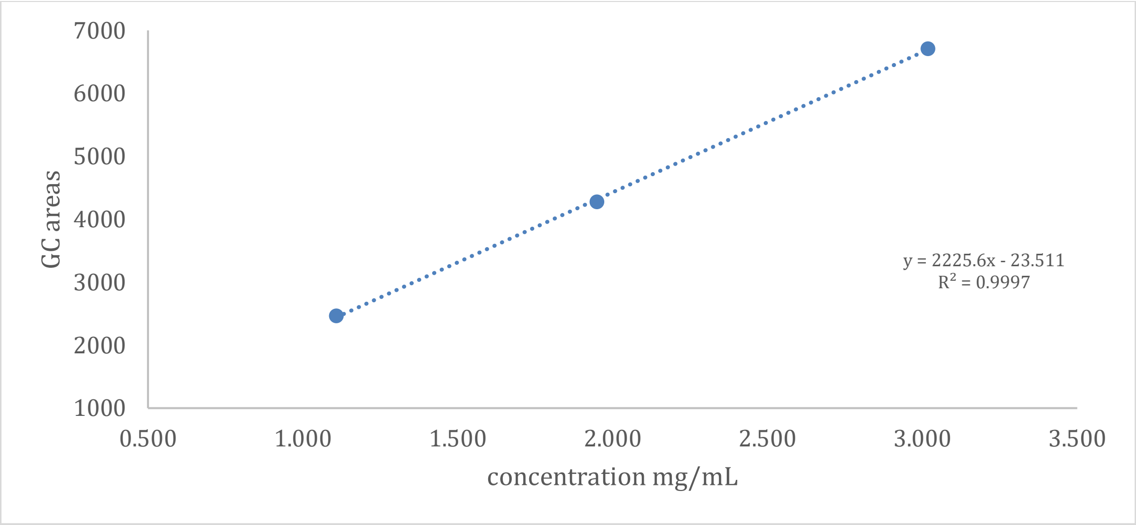


**Fig. S3.** Calibration curve for *Z*11-16Ald

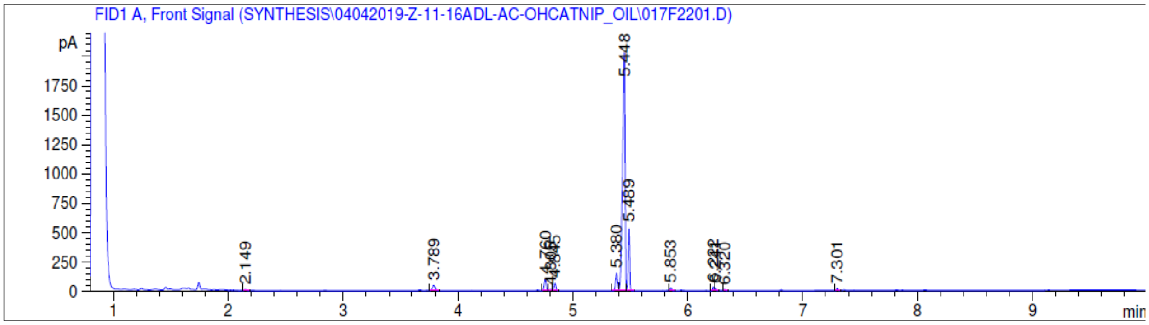


**Fig. S4.** The GC chromatogram of the biological aldehyde preparation. The peak at 5.448 min corresponds to Z11-16Ald, at 5.489 min 16Ald, and at 5.380 min Z9-16Ald. The ratio of the areas under the curve for the three C16 aldehydes was 82:13:5 (Z11-16Ald : 16Ald : Z9-16Ald).

**Table S1.** Primers used in this study

| Primer name | Primer sequence, 5’->3’ |
|---|---|
| PR-10350 (atf1_U1_fw) | agtgcagguaaaacaatgaatgaaatcgatgag |
| PR-10351 (atf1_U1_rev) | cgtgcgauctaagggcctaaaaggagagctttg |
| PR-10600 (TefYL terminator _rev) | cacgcgaUgaattcggacacgggcatc |
| PR-10655 (EpiVecYL _fw) | acccattgcUgtagatatgtcttgtgtgtaagg |
| PR-10656 (EpiVecYL _rev) | atcatgtaaUtagttatgtcacgcttacattc |
| PR-10702 (Δpex10YL_up _fw) | cattgtaactagtcctggaggg |
| PR-10703 (Δpex10YL_up _down) | acgaagttaUttgagccgaggcagatttg |
| PR-10704 (Δpex10YL_down _up) | acgaagttatUtgacgaggtctggatggaag |
| PR-10705 (Δpex10YL_down _rev) | cattgctaagaatccaaactggag |
| PR-10714 (NatMxSynYL-end_fw) | ttgcttgcgaacctaattcc |
| PR-10718 (TPex20USER _fw) | cgtgcaacgctUcagatgtgaccatatacttagg |
| PR-10719 (IntBupUSER_rev ) | aagcgttgcacgUccagtactaagactttcatgac |
| PR-10738 (Har_FAR_KKSYE_U1_rev) | cgtgcgaUttattcgtagcttttttttttccaagaaatgtctaacac |
| PR-10739 (Har_FAR_HDEL _U1_rev) | cgtgcgaUttacaattcatcatgttccaagaaatgtctaacac |
| PR-10740 (Hs_FAR_HDEL_U1_rev) | cgtgcgaUttacaattcatcatgcaagaaatgtctaacacccc |
| PR-10741 (Has_FAR_HDEL _U1_rev) | cgtgcgaUttacaattcatcatgttccaagaagtgtctaacac |
| PR-10766 (NatMxSynYL-start _fw_new) | ataacttcgUataatgtatgctatacgaagttataaggagtttggcgcccg |
| PR-10767 (NatMxSynYL-start _rev_new) | tcatggacatggcatagac |
| PR-10851 (Atrd11 expression cassette _fw) | agtgcaggUgacgcagtaggatgtcctgc |

| | |
|---|---|
| PR-10853 (Hs_Far expression cassette _fw) | acctgcacUagagaccgggttgg |
| PR-11047 (NatMxSynYL-end _rev_new) | aataacttcgUatagcatacattatacgaagttatcgagcgtcccaaaacc |
| PR-11106 (IntB_up_fw) | cgtgcgaUcccacagttctcactcag |
| PR-11107 (Hs_Far expression cassette _rev) | atgacagaUcagatgcattcttgggcgg |
| PR-11108 (URAsynYL _fw) | atctgtcaUataacttcgtataatgtatgc |
| PR-11109 (URAsynYL _rev) | cacgcgaUcataagacgcctcgttgctc |
| PR-11110 (E.coli backboneUSER _fw) | atcgcgtgcattcgcggccgcatttaaatcc |
| PR-11111 (E.coli backboneUSER _rev) | tcgcacgcattcgcggccgcaaatttaaataaaatg |
| PR-11138 (Hphsyn _fw) | agcaatgggUaaaaagcctgaactcaccgc |
| PR-11139 (Hphsyn _rev) | attacatgaUtattcctttgccctcggacg |
| PR-11446 (EpiVecYL (overhang LeuKl)_fw) | agacatUgctgtagatatgtcttgtgtgtaagg |
| PR-11447 (LeuKL (basic vector)_fw) | aatgtcUaagaatatcgttgtcctacc |
| PR-11448 (LeuKL (basic vector) _rev) | attacatgaUtaagccaagatttccttgac |
| PR-11694 (GPAT_up_USER _fw) | CGTGCGAUgcatctaggagctccattcagc |
| PR-11695 (GPAT_down _USER_rev) | CACGCGAUggacgagcagaccacg |
| PR-12989 (PrExp _fw) | CGTGCGAUaaggagtttggcgcccg |
| PR-13369 (PrExp _rev) | attggacaUtgctgtagatatgtcttg |
| PR-13370 (Cre _fw) | atgtccaaUttactgaccgtacacc |
| PR-13371 (Cre _rev) | aatcgccaUcttccagcaggcgc |
| PR-13374 (Ttef _fw) | atggcgatUagccccacgttgccggtcttg |
| PR-13494 (Nat-Tcyc-loxP_fw) | agggtacUactttggatgatactgc |

| | |
|---|---|
| PR-13549 (loxP-PrTefIntron _fw) | ataacttcgUataatgtatgctatacgaagttatagagaccgggttggcggcgc |
| PR-141 (NB326URA3fwdU) | agaacagcUgaagcttcgtacg |
| PR-14126 (Ased11_U1_fw) | agtgcaggUaaaacaatggctcaag |
| PR-14127 (Ased11_U1_rev) | cgtgcgaUttagttgtccttcc |
| PR-14128 (Sld11_U1_fw) | agtgcaggUaaaacaatggctcaat |
| PR-14129 (Sld11_U1_rev) | cgtgcgaUtcattcaccctta |
| PR-14130 (Tnd11_U1_fw) | agtgcaggUaaaacaatggctgttatg |
| PR-14131 (Tnd11_U1_rev) | cgtgcgaUtcattctttcttagcgtagaaa |
| PR-142 (NB327URA3Rev2U) | AGGCCACUAGTGGATCTGATATCAC |
| PR-14269 (UraYL_fw) | atgccctcctacgaggcccg |
| PR-14270 (UraYL_rev) | ctagcagttgatcttctggtag |
| PR-14320 (Atf1_U2_fw) | ATCTGTCAUAAAACAATGAATGAAATCGATGAG |
| PR-14321 (Atf1_U2_rev) | CACGCGAUCTAAGGGCCTAAAAGGAGAGCTTTG |
| PR-15426 (Δhfd1_up _fw) | CGTGCGAUataagaaaaaaaacag |
| PR-15427 (Δhfd1_up _rev) | AGCTGTTCUactaaccctacttcctc |
| PR-15428 (Δhfd1_down _fw) | AGTGGCCUttttattggtggtgtg |
| PR-15429 (Δhfd1_down _rev) | CACGCGAUgcatagtgcttttcatattc |
| PR-15438 (Δhfd4_up _fw) | CGTGCGAUagtatcgctactgtactaaaattg |
| PR-15439 (Δhfd4_up _rev) | AGCTGTTCUagcggacaagtgtcaatgtt |
| PR-15440 (Δhfd4_down _fw) | AGTGGCCUatgtattttatcagtagtatctc |
| PR-15441 (Δhfd4_down _rev) | CACGCGAUattggataatacatttccta |
| PR-1565 (PTEF1) | ATGACAGAUTTGTAATTAAAACTTAG |
| PR-15974 (Dmd9_U1_fw) | AGTGCAGGUAAAACAatggctccatactctagaatc |
| PR-15975 (Dmd9_U1_rev) | CGTGCGAUttatctggacttgtcaacc |

| | |
|---|---|
| PR-15976 (attB1_Dmd9_F) | GGGGACAAGTTTGTACAAAAAAGCAGGCTATGGCTCCATACTCTAGAATCTAC |
| PR-15977 (attB2_Dmd9_R) | GGGGACCACTTTGTACAAGAAAGCTGGGTTTATCTGGACTTGTCAACCAACAAAACGTTTCTAG |
| PR-15978 (attB1_Phd9_F) | GGGGACAAGTTTGTACAAAAAAGCAGGCTATGGCCCTGAAGCTGAACCCCTTC |
| PR-15979 (attB2_Phd9_R) | GGGGACCACTTTGTACAAGAAAGCTGGGTTTACAGCTTCACCTGTCGGTCGAAG |
| PR-15980 (attB1_Rcd9_F) | GGGGACAAGTTTGTACAAAAAAGCAGGCTATGGGCGTCCTGCTGAACATCTG |
| PR-15981 (attB1_Rcd9_R) | GGGGACCACTTTGTACAAGAAAGCTGGGTTTAGACCTTTCGGTCGAAGATCCA |
| PR-15982 (attB1_Atr236_F) | GGGGACAAGTTTGTACAAAAAAGCAGGCTACATGCCGCCTCAGGGTCAAGATCGCGAGTC |
| PR-15983 (att2_Atr236_R) | GGGGACCACTTTGTACAAGAAAGCTGGGTCTTATTATTCATCTTTCGAAGGGTTAAAGATGGTG |
| PR-15984 (attB1_Atr1432_F) | GGGGACAAGTTTGTACAAAAAAGCAGGCTACATGGCTCCAAATGCCACAGATGCTAATG |
| PR-15985 (attB2_Atr1432_R) | GGGGACCACTTTGTACAAGAAAGCTGGGTCTTACTAGTCATCTTTCGGGTGTATCCTTATAG |
| PR-15986 (attB1_OLE1_F) | GGGGACAAGTTTGTACAAAAAAGCAGGCTACATGCCAACTTCTGGAACTACTATTGAATTG |
| PR-15987 (attB2_OLE1_R) | GGGGACCACTTTGTACAAGAAAGCTGGGTCTTAAAAGAACTTACCAGTTTCGTAGATTTCAC |
| PR-16463 (Δfao1YL_up _fw) | CGTGCGAUTGGGGGAGGATTGCGATGGG |
| PR-16466 (Δfao1YL_down _rev) | CACGCGAUGTGTTAGTTCCTTGTAGTGTG |
| PR-16696 (GPAT_up _rev) | agctgttcUTACCGCACTTCCGGAACATC |
| PR-16698 (GPAT_100bpPr_down _fw) | agtggccUCCGATACTTGTTTGTGTGAC |
| PR-1852 (PTDH3_fw) | CACGCGAUATAAAAAACACGCTTTTTCAG |
| PR-1853 (PTDH3_rv) | ACCTGCACUTTTGTTTGTTTATGTGTGTTTATTC |
| PR-8330 (Ase_FAR_U1_fw) | agtgcagguaaaacaatgccagtcttgacttctagag |
| PR-8331 (Ase_FAR_U1_rev) | cgtgcgauttacttcttcttttcta |
| PR-8332 (Har_FAR_U1_fw) | agtgcagguaaaacaatggttgtcttgacctccaaag |
| PR-8336 (Hs_FAR_U1_fw) | agtgcagguaaaacaatggttgtcttgacctc |

| | |
|---|---|
| PR-8337 (Hs_FAR_U1_rev) | cgtgcgauttaagtcttttttcca |
| PR-8340 (Has_FAR_U1_fw) | agtgcagguaaaacaatggttgtcttgacctc |
| PR-8341 (Has_FAR_U1_rev) | cgtgcgauttacttgttggtagtct |
| PR-8350 (Atrd11_U1_fw) | agtgcagguaaaacaatggttccaaacaagggttcc |
| PR-8351 (Atrd11_U1_rev) | cgtgcgautcatctctttctacccc |
| PR-8857 (IntB A fragment _fw) | tcaggatgcacaggcacaagtacaatatatcccacagttctcactcagatc |
| PR-10604 (tracrRNA _rev) | cacgcgaUaccgtacccacacaaaaaaagcaccaccgactc |
| PR-10607 (PrtRNAGly _fw) | cgtgcgaUagtgaatcattgctaacagatc |
| PR-11694 (GPAT_up_USER _fw) | CGTGCGAUgcatctaggagctccattcagc |
| PR-11695 (GPAT_down _USER_rev) | CACGCGAUggacgagcagaccacg |
| PR-13338 (PrGPD _rev) | ACCTGCACUgttgatgtgtgtttaattc |
| PR-15607 (dsOLIGO_pex10 _KO) | aagggagtacgatttatccaccagtacaaggaggagctggagtagcgtccaagtttgcatatctcggtttgtgtacgcttgtgggctcca |
| PR-15788 (PrtRNA-Gly_rev) | taaccaaccUgcgccgacccggaatcgaac |
| PR-15789 (crRNA-TRPR_fw) | gttttagagcUagaaatagcaagttaaaataag |
| PR-15790 (gRNA_cass2_fw) | AGTGCAGGUagtgaatcattgctaacagatc |
| PR-15791 (gRNA_cass2_rev) | ACCTGCACUaccgtacccacacaaaaaaagcac |
| PR-15792 (gRNA_cass3_fw) | ATCTGTCAUagtgaatcattgctaacagatc |
| PR-15793 (gRNA_cass3_rev) | ATGACAGAUaccgtacccacacaaaaaaagcac |
| PR-15930 (PrGPDPrTefintron_rev) | acctgcggtUagtactgcaaaaagtgctgg |
| PR-16463 (Î”fao1YL_up _fw) | CGTGCGAUTGGGGGAGGATTGCGATGGG |
| PR-16465 (Î”fao1YL_down _fw) | AGTGGCCUGCAAGAGACGAGTTTAGAAATAG |

| | |
|---|---|
| PR-16466 (Î”fao1YL_down _rev) | CACGCGAUGTGTTAGTTCCTTGTAGTGTG |
| PR-16594 (Har_FAR_codop tYL_U2_fw) | aaccgcaggUGGTCCTGACCTCTAAG |
| PR-16595 (Har_FAR_codop tYL_U2_rev) | CACGCGAUCTACTCGTAGGACTTCTTCTC |
| PR-16698 (GPAT_100bpPr_ down _fw) | agtggccUCCGATACTTGTTTGTGTGAC |
| PR-17016 (gRNA1_hfd1_se nse) | CGCAGTCGCTGGATGTCGCCgttttagagct |
| PR-17017 (gRNA1_hfd1_ant isense) | GGCGACATCCAGCGACTGCGtaaccaacct |
| PR-17018 (gRNA2_hfd1_se nse) | AAGATTCGGATCAGCACGTTgttttagagct |
| PR-17019 (gRNA2_hfd1_ant isense) | AACGTGCTGATCCGAATCTTtaaccaacct |
| PR-17020 (gRNA1_hfd4_se nse) | GGATGCGTACAGGCGCCCTTgttttagagct |
| PR-17021 (gRNA1_hfd4_ant isense) | AAGGGCGCCTGTACGCATCCtaaccaacct |
| PR-17022 (gRNA2_hfd4_se nse) | GACAGAAGCGGTCCCTCCTGgttttagagct |
| PR-17023 (gRNA2_hfd4_ant isense) | CAGGAGGGACCGCTTCTGTCtaaccaacct |
| PR-17025 (dsOLIGO_hfd1_ KO) | ATTGTGTGTATATATATAATCATTCTATGAGGAAGTAGGGTTAGTTTTT ATTGGTGGTGTGTTTGAGGAAGGAGGGAGAGCGTTCAAGCT |
| PR-17026 (dsOLIGO_hfd4_ KO) | GGGTTCCTCAGACGACTTCCACGCCTTCTTCCTCACTCGCGAAGATG GAGTTTATGGGTGAGTAATGGATCGTCGCTCATGGGACTAATC |
| PR-17028 (gRNA_cass4 _fw ) | AGCTTGAGUagtgaatcattgctaacagatc |
| PR-17029 (gRNA_cass4 _rev ) | ACTCAAGCUaccgtacccacacaaaaaaagcac |
| PR-17143 (gRNA3_fao1_se nse) | TATTCGATGCCCCGGTAACTgttttagagct |
| PR-17144 (gRNA3_fao1_ant isense) | AGTTACCGGGGCATCGAATAtaaccaacct |

| | |
|---|---|
| PR-18042 (gRNA5_hfd1_sense) | GCCATGGTAAGCACCGTAACgttttagagct |
| PR-18043 (gRNA5_hfd1_antisense) | GTTACGGTGCTTACCATGGCtaaccaacct |
| PR-18107 (gRNA2_hfd2_sense) | GCCGTGATAAGACCCATAGCgttttagagct |
| PR-18108 (gRNA2_hfd2_antisense) | GCTATGGGTCTTATCACGGCtaaccaacct |
| PR-18115 (gRNA2_hfd3_sense) | GCCCCACACGATTCGCCGAGgttttagagct |
| PR-18116 (gRNA2_hfd3_antisense) | CTCGGCGAATCGTGTGGGGCtaaccaacct |
| PR-18123 (dsOLIGO_hfd2_KO) | GCCGCTTATAGCTTTGGGCGAACTGTGGTTAGTAAAAATATCATATAATCAGATAATGTGGCGATTGAACTGTATTATTTCACATGATGA |
| PR-18124 (dsOLIGO_hfd3_KO) | TACATACCCCGCTGTTATCTCTTTATCATTCATAATAAATCATATAGGCTCCAGCTTAAGGGTTAGTGGCGAGACATGTTTTCTTCAGGT |
| PR-18152 (GPAT_up (direct fusion to BB1784)_rev) | aggccacUTACCGCACTTCCGGAACATC |
| PR-18224 (fao1_up_rev_fusion) | AGGCCACUTGTCAAGTAATCAAGCTAATGC |
| PR-18913 (gRNA3_PrGPAT_sense) | CTGAGGTCTCATTTATCCAGgttttagagct |
| PR-18914 (gRNA3_PrGPAT_antisense) | CTGGATAAATGAGACCTCAGtaaccaacct |
| PR-19018 (Dmd9_U1_rev) | CGTGCGAUTTATCGAGACTTGTCC |
| PR-19102 (Dmd9_U1_fw) | AGTGCAGGUGCCACAATGGCTCCCTACTCTCG |
| PR-20733 (Fas2 (I1220F)_sense) | GGTGGTATCACCGCCCTGCGgttttagagct |
| PR-20734 (Fas2 (I1220F)_antisense) | CGCAGGGCGGTGATACCACCtaaccaacct |
| PR-20762 (Fas2 (I1220F)_up_CRISPR_repair_fw) | CCAAGTACGAGGACTACCTG |
| PR-20763 (Fas2 (I1220F)_up_CRISPR_repair_rev) | AGGGCGGUGAAACCACCCATACCGG |
| PR-20764 (Fas2_down_CRISPR_repair_fw) | ACCGCCCUGCGAGGCATGTTCAAGGACC |

| | |
|---|---|
| PR-20765 (Fas2_down_CRISPR_repair_rev) | GGAGCAGGCACAGATCGG |

**Table S2.** DNA fragments obtained by PCR using the indicated template and primers

| DNA fragment ID and name | Description | Fw_primer | Rv_primer | Template DNA |
|---|---|---|---|---|
| BB0684 | Fatty acyl-CoA reductase from *Agrotis segetum* | PR-8330 | PR-8331 | SEQ ID NO: 1 |
| BB0687 | Fatty acyl-CoA reductase from *Heliothis subflexa* | PR-8336 | PR-8337 | SEQ ID NO: 2 |
| BB0689 | Fatty acyl-CoA reductase from *Helicoverpa assulta* | PR-8340 | PR-8341 | SEQ ID NO: 3 |
| BB0694 | Δ11-desaturase from *Amyelois transitella* | PR-8350 | PR-8351 | SEQ ID NO: 4 |
| BB0914 | Fatty acyl-CoA reductase from *Helicoverpa armigera* | PR-8332 | PR-10738 | SEQ ID NO: 5 |
| BB0915 | Fatty acyl-CoA reductase from *Helicoverpa armigera* with modified C-terminus | PR-8332 | PR-10739 | SEQ ID NO: 6 |
| BB0916 | Fatty acyl-CoA reductase from *Heliothis subflexa* with modified C-terminus | PR-8336 | PR-10740 | SEQ ID NO: 7 |
| BB0917 | Fatty acyl-CoA reductase from *Helicoverpa assulta* with modified C-terminus | PR-8340 | PR-10741 | SEQ ID NO: 8 |
| BB1354 | Δ11-desaturase from *A. segetum* | PR-14126 | PR-14127 | SEQ ID NO: 9 |
| BB1355 | Δ11-desaturase from *Spodoptera littoralis* | PR-14128 | PR-14129 | SEQ ID NO: 10 |
| BB1356 | Δ11-desaturase from *Trichoplusia ni* | PR-14130 | PR-14131 | SEQ ID NO: 11 |
| BB0410 | PTDH3 promotor from *S. cerevisiae* | PR-1852 | PR-1853 | Genomic DNA of *S. cerevisiae* CEN.PK102-5B |
| BB1143 | Alcohol acetyltransferase from *S. cerevisiae* | PR-10350 | PR-10351 | Genomic DNA of *S. cerevisiae* CEN.PK102-5B |
| BB1870 | Desaturase from *Drosophila melanogaster* | PR-15976 | PR-15977 | pCfB5316 |
| BB1871 | Desaturase from *Pelargonium x hortorum* | PR-15978 | PR-15979 | pCfB4584 |
| BB1872 | Desaturase from *Ricinus communis* | PR-15980 | PR-15981 | pCfB4585 |

| | | | | |
|---|---|---|---|---|
| BB20J | Δ9-desaturase from *Amyelois transitella* | PR-15982 | PR-15983 | cDNA of *A. transitella* PG |
| BB19L | Δ9-desaturase from *Amyelois transitella* | PR-15984 | PR-15985 | cDNA of *A. transitella* PG |
| BB19J | Δ9-desaturase from *S. cerevisiae* | PR-15986 | PR-15987 | Genomic DNA of *S. cerevisiae* CEN.PK102-5B |
| BB0464 | $P_{TDH3}$ and $P_{TEF1}$ promotors from *S. cerevisiae* | PR-1565 | PR-1853 | p1977 |
| BB1696 | Desaturase from *Drosophila melanogaster* | PR-15974 | PR-15975 | SED ID NO: 12 |
| BB1422 | Alcohol acetyltransferase from *S. cerevisiae* | PR-14320 | PR-14321 | Genomic DNA of *S. cerevisiae* CEN.PK102-5B |
| BB1005 | Hygromycin resistance gene | PR-11138 | PR-11139 | SED ID NO: 13 |
| BB1006 | pCfB3405 w/o resistance gene | PR-10655 | PR-10656 | pCfB3405 w/o resistance gene |
| BB1051 | Atrd11 expression cassette and the upstream genomic region of integration site B | PR-10851 | PR-11106 | SED ID NO: 14 |
| BB1126 | Hs_FAR expression cassette | PR-10853 | PR-11107 | SEQ ID NO: 15 |
| BB1131 | Hs_FAR expression cassette | PR-10853 | PR-10655 | pCfB3465 |
| BB1132 | Part of pCfB3465 vector | PR-10656 | PR-10851 | pCfB3465 |
| BB1135 | Vector backbone for propagation in *E.coli* | PR-11110 | PR-11111 | pCfB2196 |
| BB1137 | Ura3 marker cassette fused to 500 bp downstream region for integration into IntB site | PR-11108 | PR-11109 | SEQ ID NO: 16 |
| BB1338 | Hygromycin resistance marker | PR-141 | PR-142 | pCfB6574 |
| BB1346 | Nourseothricin resistance marker | PR-141 | PR-142 | pCfB4848 |
| BB1349 | Genomic region upstream of *pex10* fused to 2/3 Start of nourseothricin resistance cassette | PR-10702 | PR-10767 | BB1144/BB1347 |
| BB1350 | Genomic region downstream of *pex10* fused to 2/3 end of nourseothricin resistance cassette | PR-13494 | PR-10705 | BB1348/BB1145 |
| BB1427 | Ura3 marker cassette | PR-141 | PR-142 | pCfB4586 |

| | | | | |
|---|---|---|---|---|
| BB1543 | Genomic region upstream of *hfd1* | PR-15426 | PR-15427 | Genomic DNA *Yarrowia lipolytica* GB20 |
| BB1544 | Genomic region downstream of *hfd1* | PR-15428 | PR-15429 | Genomic DNA *Yarrowia lipolytica* GB20 |
| BB1549 | Genomic region upstream of *hfd4* | PR-15438 | PR-15439 | Genomic DNA *Yarrowia lipolytica* GB20 |
| BB1550 | Genomic region downstream of *hfd4* | PR-15440 | PR-15441 | Genomic DNA *Yarrowia lipolytica* GB20 |
| BB1757 | Genomic region upstream of *fao1* fused to 2/3 start of Ura3 cassette | PR-16463 | PR-14270 | BB1725/BB1427 |
| BB1758 | Genomic region downstream of *fao1* fused to 2/3 end of Ura3 cassette | PR-14269 | PR-16466 | BB1726/BB1427 |
| BB1782 | Genomic region upstream of *pex10* | PR-11694 | PR-16696 | Genomic DNA *Yarrowia lipolytica* GB20 |
| BB1784 | Genomic region downstream of *pex10* | PR-16698 | PR-11695 | Genomic DNA *Yarrowia lipolytica* GB20 |
| BB1144 | Genomic region upstream of *pex10* | PR-10702 | PR-10703 | Genomic DNA *Yarrowia lipolytica* GB20 |
| BB1347 | 2/3 Start of nourseothricin resistance cassette | PR-13549 | PR-10767 | pCfB4848 |
| BB1348 | 2/3 End of nourseothricin resistance cassette | PR-13494 | PR-11047 | pCfB4848 |
| BB1145 | Genomic region downstream of *pex10* | PR-10704 | PR-10705 | Genomic DNA *Yarrowia lipolytica* GB20 |
| BB1048 | Integration site B | PR-8857 | PR-10719 | Genomic DNA *Yarrowia lipolytica* GB20 |
| BB1050 | Atrd11 expression cassette | PR-10851 | PR-10718 | SEQ ID NO: 17 |
| BB1352 | Genomic region upstream of *pex10* fused to 2/3 Start of nourseothricin resistance cassette | PR-10702 | PR-10767 | BB1144/BB1014 |

| | | | | |
|---|---|---|---|---|
| BB1353 | Genomic region downstream of *pex10* fused to 2/3 end of nourseothricin resistance cassette | PR-10714 | PR-10705 | BB1351/BB1145 |
| BB1288 | EXP promoter | PR-12989 | PR-13369 | pCfB3405 |
| BB1289 | Cre recombinase | PR-13370 | PR-13371 | pSH66 |
| BB1291 | TEF1 terminator | PR-13374 | PR-10600 | Genomic DNA *Yarrowia lipolytica* GB20 |
| BB1129 | LEU2 from Kluyveromyces lactis | PR-11447 | PR-11448 | SEQ ID NO: 18 |
| BB1130 | pCfB3431 w/o resistance gene | PR-11446 | PR-10656 | pCfB3431 |
| BB1014 | 2/3 Start of nourseothricin resistance cassette | PR-10766 | PR-10767 | pCfB3405 |
| BB1351 | 2/3 End of nourseothricin resistance cassette | PR-10714 | PR-11047 | pCfB3405 |
| BB1726 | Upstream region of *fao1* | PR-16463 | PR-18224 | Genomic DNA *Yarrowia lipolytica* GB20 |
| BB1784 | Genomic DNA downstream of GPAT promoter | PR-16698 | PR-11695 | Genomic DNA *Yarrowia lipolytica* GB20 |
| BB2081 | Genomic DNA upstream of GPAT promoter | PR-11694 | PR-18152 | Genomic DNA *Yarrowia lipolytica* GB20 |
| BB2102 | Downstream region of *fao1* | PR-16465 | PR-16466 | Genomic DNA *Yarrowia lipolytica* GB20 |
| BB2103 | Up- and downstream genomic region around *fao1* | PR-16463 | PR-16466 | BB2102/BB1726 |
| BB2082 | Up- and downstream genomic region GPAT promoter | PR-11694 | PR-11695 | BB1784/BB2081 |
| BB2311 | Upstream genomic region of $Fas2p^{I1220}$ | PR-20762 | PR-20763 | Genomic DNA *Yarrowia lipolytica* GB20 |
| BB2312 | Downstream genomic region of $Fas2p^{I1220}$ | PR-20764 | PR-20765 | Genomic DNA *Yarrowia lipolytica* GB20 |
| BB2313 | Up- and downstream genomic region of $Fas2p^{I1220}$ | PR-20762 | PR-20765 | BB2311/BB2312 |
| BB1892 | gRNA expression cassette targeting *hfd1* | PR-10607 | PR-15791 | BB1635, BB1636, PR-17016, PR-17017 |
| BB1893 | gRNA expression cassette targeting *hfd1* | PR-15790 | PR-15793 | BB1635, BB1636, PR-17018, PR-17019 |

| | | | | |
|---|---|---|---|---|
| BB1894 | gRNA expression cassette targeting *hfd4* | PR-15792 | PR-17029 | BB1635, BB1636, PR-17020, PR-17021 |
| BB1895 | gRNA expression cassette targeting *hfd4* | PR-17028 | PR-10604 | BB1635, BB1636, PR-17022, PR-17023 |
| BB1635 | Promoter for gRNA expression | PR-10607 | PR-15788 | pCfB4589 |
| BB1636 | Terminator region for gRNA expression | PR-15789 | PR-10604 | pCfB4589 |
| BB1687 | Fused promoter regions of *gpd and tef1* | PR-13338 | PR-15930 | pCfB3465 |
| BB1740 | HarFAR reductase | PR-16594 | PR-16595 | SEQ ID NO: 19 |
| BB2206 | Dmd9 desaturase | PR-19102 | PR-19018 | SEQ ID NO: 20 |

**Table S3.** Plasmids used in this study

| Vector name | Selection marker for yeast | Parent vector | BioBricks | Reference |
|---|---|---|---|---|
| pCfB2190 | KlLEU2 | - | - | (3) |
| pCfB2228 | SpHIS5 | - | - | (3) |
| pCfB2501 | KlLEU2 | pCfB2190 | BB0410, BB0684 | This study |
| pCfB2504 | KlLEU2 | pCfB2190 | BB0410, BB0687 | This study |
| pCfB2506 | KlLEU2 | pCfB2190 | BB0410, BB0689 | This study |
| pCfB2537 | SpHIS5 | pCfB2228 | BB0410, BB0694 | This study |
| pCfB3412 | KlLEU2 | pCfB2190 | BB0410, BB0914 | This study |
| pCfB3413 | KlLEU2 | pCfB2190 | BB0410, BB0915 | This study |
| pCfB3414 | KlLEU2 | pCfB2190 | BB0410, BB0916 | This study |
| pCfB3415 | KlLEU2 | pCfB2190 | BB0410, BB0917 | This study |
| pCfB4369 | SpHIS5 | pCfB2228 | BB0410, BB1354 | This study |
| pCfB4370 | SpHIS5 | pCfB2228 | BB0410, BB1355 | This study |
| pCfB4371 | SpHIS5 | pCfB2228 | BB0410, BB1356 | This study |
| pYEX-CHT-DEST | Ura3 | - | - | (7) |
| pYEX-CHT-Atrd1432 | Ura3 | pYEX-CHT-DEST | BB19L | This study |
| pYEX-CHT-Atr236 | Ura3 | pYEX-CHT-DEST | BB20J | This study |
| pYEX-CHT-Phd9 | Ura3 | pCfB4584 | BB1871 | This study |
| pYEX-CHT-Rcd9 | Ura3 | pCfB4585 | BB1872 | This study |
| pYEX-CHT-OLE1 | Ura3 | pYEX-CHT-DEST | BB19J | This study |
| pYEX-CHT-Dmd9 | Ura3 | pCfB5316 | BB1870 | This study |
| pCfB2909 | None | - | - | (4) |
| p1977 | None | - | - | (4) |
| pCfB4580 | KlLEU2 | pCfB2190 | BB0464, BB0915, BB1422 | This study |

| | | | | |
|---|---|---|---|---|
| pCfB5316 | None | pCfB2909 | BB0410, BB1696 | This study |
| pCfB3465 | Ura3 | - | BB1051, BB1126, BB1135, BB1137 | This study |
| pCfB5110 | NatSyn | - | BB1135, BB1346, BB1543, BB1544 | This study |
| pCfB5113 | HphSyn | - | BB1135, BB1338, BB1549, BB1550 | This study |
| pCfB3516 | HphSyn | - | BB1005, BB1131, BB1132 | This study |
| pCfB5573 | HphSyn | - | BB1135, BB1338, BB1757, BB1758 | This study |
| pCfB5750 | Ura3 | - | BB1135, BB1427, BB1782, BB1784 | This study |
| pCfB2196 | BleMX | - | - | (3) |
| pCfB4158 | KlLEU2 | pCfB3529 | BB1288, BB1289, BB1291 | This study |
| pCfB3529 | KlLEU2 | - | BB1129, BB1130 | This study |
| pCfB3431 | HphSyn | - | BB1006, BB1005 | This study |
| pCfB3405 | NatSyn | - | - | (2) |
| pCfB6574 | HphSyn | - | - | (2) |
| pCfB4848 | NatSyn | - | - | (2) |
| pCfB4586 | Ura3 | - | - | (2) |
| pSH66 | - | - | - | EuroScarf |
| pCfB6364 | - | - | - | (8) |
| pCfB3405 | - | - | - | (2) |
| pCfB5878 | NatSyn | pCfB3405 | BB1892, BB1893, BB1894, BB1895 | |
| pCfB6530 | NatSyn | pCfB3405 | BB1635, BB1636, PR-18042, PR-18043 | |
| pCfB5790 | NatSyn | pCfB3405 | BB1833, BB1834 | |
| pCfB6463 | NatSyn | pCfB3405 | BB1635, BB1636, PR-17143, PR-17144 | |
| pCfB6566 | NatSyn | pCfB3405 | BB1635, BB1636, PR-18107, PR-18108 | |
| pCfB6570 | NatSyn | pCfB3405 | BB1635, BB1636, PR-18115, PR-18116 | |
| pCfB6834 | NatSyn | pCfB3405 | BB1635, BB1636, PR-18913, PR-18914 | |
| pCfB6627 | - | - | - | (2) |
| pCfB6682 | - | - | - | (2) |
| pCfB6969 | None | pCfB6682 | BB1687, BB1740, BB2206 | |

| | | | | |
|---|---|---|---|---|
| pCfB7088 | NatSyn | pCfB3405 | BB1635, BB1636, PR-20733, PR-20734 | |
| pCfB4589 | - | - | - | (2) |

**Table S4.** Yeast strains

| Strain name | Strain description | Parent strain | Plasmids/BioBricks integrated | Reference |
|---|---|---|---|---|
| *Saccharomyces cerevisiae* CEN.PK102-5B | MATa *ura3-52 his3□1 leu2-3/112 MAL2-8c SUC2* | - | - | |
| ST3706 | Atrd11, Har_FAR_HDEL | CEN.PK102-5B | pCfB2537, pCfB3413 | This study |
| ST4487 | Ased11, Har_FAR_HDEL | CEN.PK102-5B | pCfB4369, pCfB3413 | This study |
| ST4488 | Sld11, Har_FAR_HDEL | CEN.PK102-5B | pCfB4370, pCfB3413 | This study |
| ST4489 | Tnd11, Har_FAR_HDEL | CEN.PK102-5B | pCfB4371, pCfB3413 | This study |
| ST3328 | Atrd11, Ase_FAR | CEN.PK102-5B | pCfB2537, pCfB2501 | This study |
| ST3339 | Atrd11, Has_FAR | CEN.PK102-5B | pCfB2537, pCfB2506 | This study |
| ST3708 | Atrd11, Has_FAR_HDEL | CEN.PK102-5B | pCfB2537, pCfB3415 | This study |
| ST3330 | Atrd11, Hs_FAR | CEN.PK102-5B | pCfB2537, pCfB2504 | This study |
| ST3707 | Atrd11, Hs_FAR_HDEL | CEN.PK102-5B | pCfB2537, pCfB3414 | This study |
| ST3705 | Atrd11, Har_FAR | CEN.PK102-5B | pCfB2537, pCfB3412 | This study |
| *S. cerevisiae* W303a *Δelo1Δole1* | MATa *elo1::HIS3 ole1::LEU2 ade2 his3 leu2 ura3* | - | - | (9) |
| ST_Atr1432 | Atr1432 | W303a *Δelo1Δole1* | pYEX-CHT-Atrd1432 | This study |
| ST_Atr236 | Atr236 | W303a *Δelo1Δole1* | pYEX-CHT-Atr236 | This study |
| ST_Phd9 | Phd9 | W303a *Δelo1Δole1* | pYEX-CHT-Phd9 | This study |
| ST_Rcd9 | Rcd9 | W303a *Δelo1Δole1* | pYEX-CHT-Rcd9 | This study |
| ST_ScOLE1 | ScOLE1 | W303a *Δelo1Δole1* | pYEX-CHT-OLE1 | This study |
| ST_DmeΔ9 | DmeΔ9 | W303a *Δelo1Δole1* | pYEX-CHT-Dmd9 | This study |
| ST4854 | ScATF1, Har_FAR | CEN.PK102-5B | pCfB4580 | This study |
| ST5290 | ScATF1, Har_FAR, DmeΔ9 | CEN.PK102-5B | pCfB4580, pCfB5316 | This study |
| *Yarrowia lipolytica* GB20 | *mus51Δ*, nugm-Htg2, ndh2i, lys11$^-$, leu2$^-$, ura3$^-$, MatB | - | - | (10) |
| ST3844 | IntB::Atrd11 Hs_FAR | *Y. lipolytica* GB20 | pCfB3465 | This study |
| ST5107 | IntB::Atrd11 Hs_FAR *Δhfd1* | ST3844 | pCfB5110 | This study |

| | | | | |
|---|---|---|---|---|
| ST5110 | IntB::Atrd11 Hs_FAR *Δhfd4* | ST3844 | pCfB5113 | This study |
| ST3842 | IntB::Atrd11 Hs_FAR *Δpex10* | ST3737 | pCfB3516 | This study |
| ST5255 | IntB::Atrd11 Hs_FAR *Δhfd1 Δhfd4* | ST5107 | pCfB5113 | This study |
| ST5452 | IntB::Atrd11 Hs_FAR *Δhfd1 Δhfd4 Δpex10* | ST5450 | BB1349/BB1350 | This study |
| ST5789 | IntB::Atrd11 Hs_FAR *Δhfd1 Δhfd4 Δpex10 Δfao1* | ST5452 | pCfB5573 | This study |
| ST5791 | IntB::Atrd11 Hs_FAR *Δhfd1 Δhfd4 Δpex10 Δfao1* ΔPrGPAT | ST5789 | pCfB5750 | This study |
| ST6289 | IntB::Atrd11 Hs_FAR *Δhfd1 Δhfd4 Δpex10 Δfao1* ΔPrGPAT IntE4::Atrd11 HarFAR | ST5790 | pCfB5751 | This study |
| ST6379 | IntB::Atrd11 Hs_FAR *Δhfd1 Δhfd4 Δpex10 Δfao1* ΔPrGPAT IntE4::Atrd11 HarFAR IntC1::Atrd11 HarFAR | ST5791 | pCfB5752 | This study |
| ST3737 | Δ*pex10* | *Y. lipolytica* GB20 | BB1352/BB1353 | This study |
| ST5449 | IntB::Atrd11 Hs_FAR *Δhfd1 Δhfd4* pCfB4158 | ST5452 | pCfB4158 (replicative) | This study |
| ST5450 | IntB::Atrd11 Hs_FAR *Δhfd1 Δhfd4* | ST5449 | pCfB4158 (replicative) lost | This study |
| ST4840 (Y-17536) | *Wild-type Yarrowia lipolytica* | | | Agricultural Research Service (NRRL, USA) |
| ST6029 | *ku70*Δ Cas9 | ST4840 | pCfB6364 | This study |
| ST6166 | *ku70*Δ Cas9 *hfd4*Δ | ST6029 | PCfB5878/PR-17026 | This study |
| ST6278 | *ku70*Δ Cas9 *hfd4*Δ *hfd1*Δ | ST6160 | pCfB6530/PR-17025 | This study |
| ST6285 | *ku70*Δ Cas9 *hfd4*Δ *hfd1*Δ *pex10*Δ | ST6278 | pCfB5790/PR-15607 | This study |
| ST6309 | *ku70*Δ Cas9 *hfd4*Δ *hfd1*Δ *pex10*Δ *fao1*Δ | ST6285 | pCfB6463/BB2103 | This study |
| ST6526 | *ku70*Δ Cas9 *hfd4*Δ *hfd1*Δ *pex10*Δ *fao1*Δ *hfd2*Δ | ST6309 | PCfB6566/PR-18123 | This study |
| ST6541 | *ku70*Δ Cas9 *hfd4*Δ *hfd1*Δ *pex10*Δ *fao1*Δ *hfd2*Δ *hfd3*Δ | ST6526 | pCfB6570/PR-18124 | This study |
| ST6629 | *ku70*Δ Cas9 *hfd4*Δ *hfd1*Δ *pex10*Δ *fao1*Δ *hfd2*Δ *hfd3*Δ *GPAT_100bpPr* | ST6541 | pCfB6834/BB2082 | This study |

| | | | | |
|---|---|---|---|---|
| ST6713 | *ku70*Δ Cas9 *hfd4*Δ *hfd1*Δ *pex10*Δ *fao1*Δ *hfd2*Δ *hfd3*Δ *GPAT_100bpPr* IntC_2:Dmd9_HarFAR | ST6629 | pCfB6627/pCfB6969 | This study |
| ST7010 | *ku70*Δ Cas9 *hfd4*Δ *hfd1*Δ *pex10*Δ *fao1*Δ *hfd2*Δ *hfd3*Δ *GPAT_100bpPr* IntC_2:Dmd9_HarFAR Fas2p$^{I1220F}$ | ST6713 | pCfB7088/BB2313 | This study |

**Table S5.** Strain genotypes on figures

| | |
|---|---|
| **Fig. 1b.** The following strains are shown in the order left to right: | |
| | *Saccharomyces cerevisiae* CEN.PK102-5B (MATa *ura3-52 his3Δ1 leu2-3/112 MAL2-8c SUC2*) |
| ST3706 | *S. cerevisiae* CEN.PK102-5B Atrd11, Har_FAR_HDEL |
| ST4487 | *S. cerevisiae* CEN.PK102-5B Ased11, Har_FAR_HDEL |
| ST4488 | *S. cerevisiae* CEN.PK102-5B Sld11, Har_FAR_HDEL |
| ST4489 | *S. cerevisiae* CEN.PK102-5B Tnd11, Har_FAR_HDEL |
| ST3328 | *S. cerevisiae* CEN.PK102-5B Atrd11, Ase_FAR |
| ST3339 | *S. cerevisiae* CEN.PK102-5B Atrd11, Has_FAR |
| ST3708 | *S. cerevisiae* CEN.PK102-5B Atrd11, Has_FAR_HDEL |
| ST3330 | *S. cerevisiae* CEN.PK102-5B Atrd11, Hs_FAR |
| ST3707 | *S. cerevisiae* CEN.PK102-5B Atrd11, Hs_FAR_HDEL |
| ST3705 | *S. cerevisiae* CEN.PK102-5B Atrd11, Har_FAR |
| **Fig. 1d.** The following strains are shown in the order left to right: | |
| | *S. cerevisiae* W303a *Δelo1Δole1* (*MATa elo1::HIS3 ole1::LEU2 ade2 his3 leu2 ura3*) |
| ST_Atr1432 | *S. cerevisiae* W303a *Δelo1Δole1*, Atr1432 |
| ST_Atr236 | *S. cerevisiae* W303a *Δelo1Δole1*, Atr236 |
| ST_Phd9 | *S. cerevisiae* W303a *Δelo1Δole1*, Phd9 |
| ST_Rcd9 | *S. cerevisiae* W303a *Δelo1Δole1*, Rcd9 |
| ST_ScOLE1 | *S. cerevisiae* W303a *Δelo1Δole1*, ScOLE1 |
| ST_DmeD9 | *S. cerevisiae* W303a *Δelo1Δole1*, DmeD9 |
| **Fig. 1e.** The following strains are shown in the order left to right: | |
| ST4854 | *S. cerevisiae* CEN.PK102-5B ScATF1, Har_FAR |
| ST5290 | *S. cerevisiae* CEN.PK102-5B ScATF1, Har_FAR, DmeΔ9 |
| **Fig. 2a.** The following strains are shown in the order left to right: | |
| | *Yarrowia lipolytica* GB20 (*mus51Δ*, nugm-Htg2, ndh2i, lys11$^{-}$, leu2$^{-}$, ura3$^{-}$, MatB) |
| ST3844 | *Y. lipolytica* GB20 IntB::Atrd11 Hs_FAR |
| ST5107 | *Y. lipolytica* GB20 IntB::Atrd11 Hs_FAR, *hfd1Δ* |
| ST5110 | *Y. lipolytica* GB20 IntB::Atrd11 Hs_FAR, *hfd4Δ* |
| ST3842 | *Y. lipolytica* GB20 IntB::Atrd11 Hs_FAR, *pex10Δ* |
| ST5255 | *Y. lipolytica* GB20 IntB::Atrd11 Hs_FAR, *hfd1Δ, hfd4Δ* |
| ST5452 | *Y. lipolytica* GB20 IntB::Atrd11 Hs_FAR, *hfd1Δ, hfd4Δ, pex10Δ* |
| ST5789 | *Y. lipolytica* GB20 IntB::Atrd11 Hs_FAR, *hfd1Δ, hfd4Δ, pex10Δ, fao1Δ* |
| ST5791 | *Y. lipolytica* GB20 IntB::Atrd11 Hs_FAR, *hfd1Δ, hfd4Δ, pex10Δ,fao1Δ,* $P_{GPAT}$*::100* |
| **Fig. 3a.** The following strains are shown in the order left to right: | |
| ST5791 | *Y. lipolytica* GB20 IntB::Atrd11 Hs_FAR, *hfd1Δ, hfd4Δ, pex10Δ,fao1Δ,* $P_{GPAT}$*::100* |
| ST6289 | *Y. lipolytica* GB20 IntB::Atrd11 Hs_FAR, *hfd1Δ, hfd4Δ, pex10Δ,fao1Δ,* $P_{GPAT}$*::100*, IntE4::Atrd11 HarFAR |
| ST6379 | *Y. lipolytica* GB20 IntB::Atrd11 Hs_FAR, *hfd1Δ, hfd4Δ, pex10Δ,fao1Δ,* $P_{GPAT}$*::100*, IntE4::Atrd11 HarFAR, IntC1::Atrd11 HarFAR |
| **Fig. 3b.** The following strains are shown in the order left to right: | |
| ST6713 | *ku70Δ* Cas9 *hfd4Δ hfd1Δ pex10Δ fao1Δ hfd2Δ hfd3Δ GPAT_100bpPr* IntC_2:Dmd9_HarFAR |
| ST7010 | *ku70Δ* Cas9 *hfd4Δ hfd1Δ pex10Δ fao1Δ hfd2Δ hfd3Δ GPAT_100bpPr* IntC_2:Dmd9_HarFAR Fas2p$^{I1220F}$ |

**Table S6.** Quantification of Z11-16:Ald

| | Sample size (µl) | Volume of flask (ml) | Area GC | Concentration of Z11-16-Ald in diluted sample (mg/ml) | Concentration Z11-16-Ald (mg/ml) |
|---|---|---|---|---|---|
| Replicate 1 | 200 | 5 | 3147.6 | 1.425 | 35.62 |
| Replicate 2 | 200 | 5 | 3106.6 | 1.406 | 35.16 |
| Replicate 3 | 200 | 5 | 3109.9 | 1.408 | 35.20 |

**SI References**


1. N. B. Jensen, *et al.*, EasyClone: method for iterative chromosomal integration of multiple genes in *Saccharomyces cerevisiae*. *FEMS Yeast Res.* **14**, 238–248 (2014).

2. C. Holkenbrink, *et al.*, EasyCloneYALI: CRISPR/Cas9-based synthetic toolbox for engineering of the yeast *Yarrowia lipolytica*. *Biotechnology Journal* **13**, 1700543 (2018).

3. V. Stovicek, I. Borodina, J. Forster, CRISPR–Cas system enables fast and simple genome editing of industrial *Saccharomyces cerevisiae* strains. *Metabolic Engineering Communications* **2**, 13–22 (2015).

4. M. M. Jessop-Fabre, *et al.*, EasyClone-MarkerFree: A vector toolkit for marker-less integration of genes into *Saccharomyces cerevisiae* via CRISPR-Cas9. *Biotechnology Journal* **11**, 1110–1117 (2016).

5. J. Dahlin, *et al.*, Multi-omics analysis of fatty alcohol production in engineered yeasts *Saccharomyces cerevisiae* and *Yarrowia lipolytica*. *Front. Genet.* **10** (2019).

6. S. Kikionis, E. Ioannou, M. Konstantopoulou, V. Roussis, Electrospun Micro/Nanofibers as Controlled Release Systems for Pheromones of Bactrocera oleae and Prays oleae. *J. Chem. Ecol.* **43**, 254–262 (2017).

7. B.-J. Ding, C. Carraher, C. Löfstedt, Sequence variation determining stereochemistry of a Δ11 desaturase active in moth sex pheromone biosynthesis. *Insect Biochem. Mol. Biol.* **74**, 68–75 (2016).

8. E. R. Marella, *et al.*, A single-host fermentation process for the production of flavor lactones from non-hydroxylated fatty acids. *Metabolic Engineering* (2019) https:/doi.org/10.1016/j.ymben.2019.08.009 (June 12, 2020).

9. R. Schneiter, V. Tatzer, G. Gogg, E. Leitner, S. D. Kohlwein, Elo1p-dependent carboxy-terminal elongation of C14:1Δ9 to C16:1Δ11 fatty acids in *Saccharomyces cerevisiae*. *J Bacteriol* **182**, 3655–3660 (2000).

10. H. Angerer, *et al.*, The LYR protein subunit NB4M/NDUFA6 of mitochondrial complex I anchors an acyl carrier protein and is essential for catalytic activity. *Proc Natl Acad Sci U S A* **111**, 5207–5212 (2014).
